# Cannabis-enriched oral *Actinomyces* induces anxiety-like behavior via impairing mitochondria and GABA signaling

**DOI:** 10.1101/2025.11.21.689724

**Authors:** Tabinda Salman, Zhenwu Luo, Douglas Johnson, Arshad A. Noorani, Zhuang Wan, Bogdan Bordieanu, Zhi-wei Ye, Rachel D. Penrod, Hongxu Xian, Sylvia Fitting, Peter W. Kalivas, Wei Jiang

## Abstract

The human oral microbiome is increasingly recognized as a contributor to brain health, yet its mechanisms remain unclear. Our previous work revealed that oral *Actinomyces* species was enriched in chronic cannabis smokers. Here, we show oral inoculation of cannabis use-associated *Actinomyces* species, especially *A. meyeri*, to wild-type C57BL/6 mice leads to anxiety-like behaviors, non-region-specific microglia activation, mitochondrial dysfunction, and reduced GABAergic neurotransmission, without evidence of bacterial translocation to the brain, neuroinflammation, and memory decline. Notably, *Actinomyces* species-producing metabolites, i.e., arginine and argininosuccinate, were increased in both oral swabs and brain following inoculation *in vivo*. These *Actinomyces* species-producing metabolites induced mitochondrial dysfunction and oxidative stress in neurons *in vitro*, indicating a neuropathogenic role and aligning with reduced GABAergic neurotransmission in vivo. Together, these results suggest that oral cannabis-associated dysbiosis impacts behavior through mitochondrial stress and impaired inhibitory signaling, indicating the oral–brain metabolic axis is potentially consequential in neuropsychiatric disorders.

**Teaser:** Chronic heavy cannabis use-enriched oral bacteria can drive anxiety and neuropathogenesis in mice.

**Highlights:**

◦ Cannabis-associated oral Actinomyces enrichment induces anxiety-like behavior in mice
◦ Microglial activation occurs without neuroinflammation (IL-1β, TNF-α, and IL-6)
◦ Mitochondrial hyperactivation and reduced inhibitory GABAergic signaling

**Graphical Abstract:** 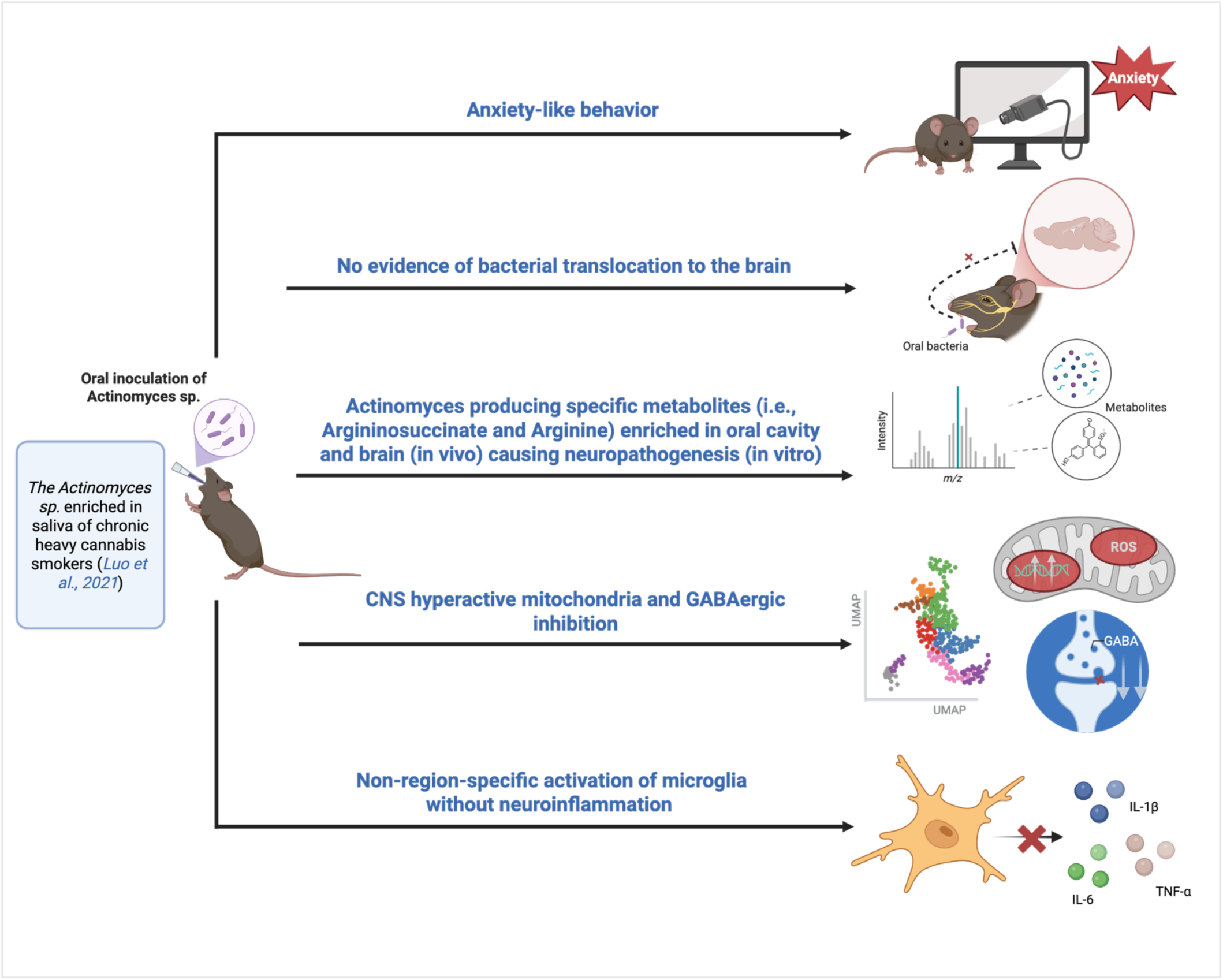

## INTRODUCTION

Cannabis is widely used around the world for medicinal and recreational purposes. The rising prevalence of cannabis use in adolescence is well documented (*1, 2*) and is accompanied by a declining perception of risk among the general population (*3, 4*). The increasing legalization of marijuana, along with earlier age of onset, higher frequency of use, and greater overall cannabis exposure, are concerning trends that represent significant risk factors for a variety of chronic cannabis use-related disorders (*1, 3*). Acute or occasional, edible, and cannabidiol (CBD)-enriched cannabis use may reduce anxiety (*5, 6*). Increasing studies indicate that chronic use of cannabis and its primary psychoactive component, tetrahydrocannabinol (THC), is associated with a range of psychopathological illnesses (*4, 7*). Young individuals who consume high-THC cannabis exhibit greater vulnerability to developing anxiety and cannabis use disorder (*8*), other psychiatric-like symptoms (*9, 10*), and frequent relapses (*11*). Therefore, it is critical that clinical and preclinical investigations of chronic cannabinoid exposure, particularly during adolescence, address its potential to remodel the brain and drive long-lasting behavioral changes later in life.

The oral microbiota is reported to be influenced by smoking and linked with various pathophysiological conditions (*12, 13*). Cannabis/cannabinoid use can have a positive or negative impact on the oral microbiome, as well as being associated with a variety of disorders, including cognitive deficiency, anxiety/depression, inflammation, oral disease, or obesity (*14*). We previously reported that chronic heavy cannabis smokers showed a disrupted balance of bacteria (specifically enriched *Actinomyces* genus and species) in their mouth (*15*). Although the *Actinomyces* genus is considered predominantly commensal in the human oral cavity (*16*), *Actinomyces* species, such as *A. meyeri,* can act as opportunistic pathogens (*17*). Building on the findings of Luo et al. 2021, (*15*), which showed enrichment of oral *Actinomyces* in chronic heavy cannabis smokers, and given that *Actinomyces* are generally reported to be less prominent in the gut due to environmental constraints, we focused on *Actinomyces* as a representative microbiome candidate of cannabis-associated oral dysbiosis.

Anxiety disorders are among the most prevalent neuropsychiatric conditions globally, yet the underlying biological mechanisms are often poorly understood (*18*). Genetics (*19*), immune dysregulations (*20*), and central nervous system (CNS) factors like neurotransmitter imbalances, particularly reduced inhibitory GABAergic signaling, have been implicated in anxiety-related pathophysiology (*21*). Additionally, peripheral influences such as the gut-brain and oral-brain axes are increasingly identified as potential modulators of emotional regulation (*22, 23*). Chronic cannabis use has been associated with both heightened anxiety and alterations in the oral microbiome in previous human studies (*15, 24*). Thus, to investigate the causality of cannabis-associated oral microbiomes on anxiety and neuropathogenesis, we orally inoculated B6 mice and evaluated neurobehavioral alterations, such as anxiety-like behavioral symptoms and disrupted inhibitory neurotransmission and mitochondrial function.

## RESULTS

### Chronic oral *A. meyeri* inoculation induces anxiety-like behavior

Following five days of oral antibiotic treatment to support the target microbiome colonization (Fig. 1A), two cannabis use-enriched *Actinomyces* species, *A. meyeri* (bacteria of interest) and *A. odontolyticus*, as well as a cannabis use-depleted species, *N. elongata* (*15*), and a vehicle control, were orally inoculated to wild-type C57BL/6 (B6) mice for six months. Only animals with confirmed colonization of the inoculated bacteria were included in the analyses. After confirming successful oral colonization of the bacteria of interest (Fig. S1), mice were subjected to different behavioral tests (baseline locomotor activity, weight, and serum corticosterone levels following six months of inoculation), before the battery of mouse behavioral tests (Fig. 1A and Fig. S2A). After six months of oral inoculation of *A. meyeri*, mice spent more time in the periphery of the open field test and less time in the absolute center of the apparatus as compared to the vehicle control, suggesting anxiety-like open field behavior (Fig. 1B and C). The two *Actinomyces* species-treated mice, when subjected to the elevated plus-maze test, avoided the open arm of the apparatus and preferred the closed arm as compared to the vehicle control, suggesting that these mice exhibited anxiety-like behavior (Fig. 1D and E). There was no significant alteration in cognition or depression related behaviors in *Actinomyces* species-treated mice (Supplementary Fig. S2B-D).

**Fig. 1:**
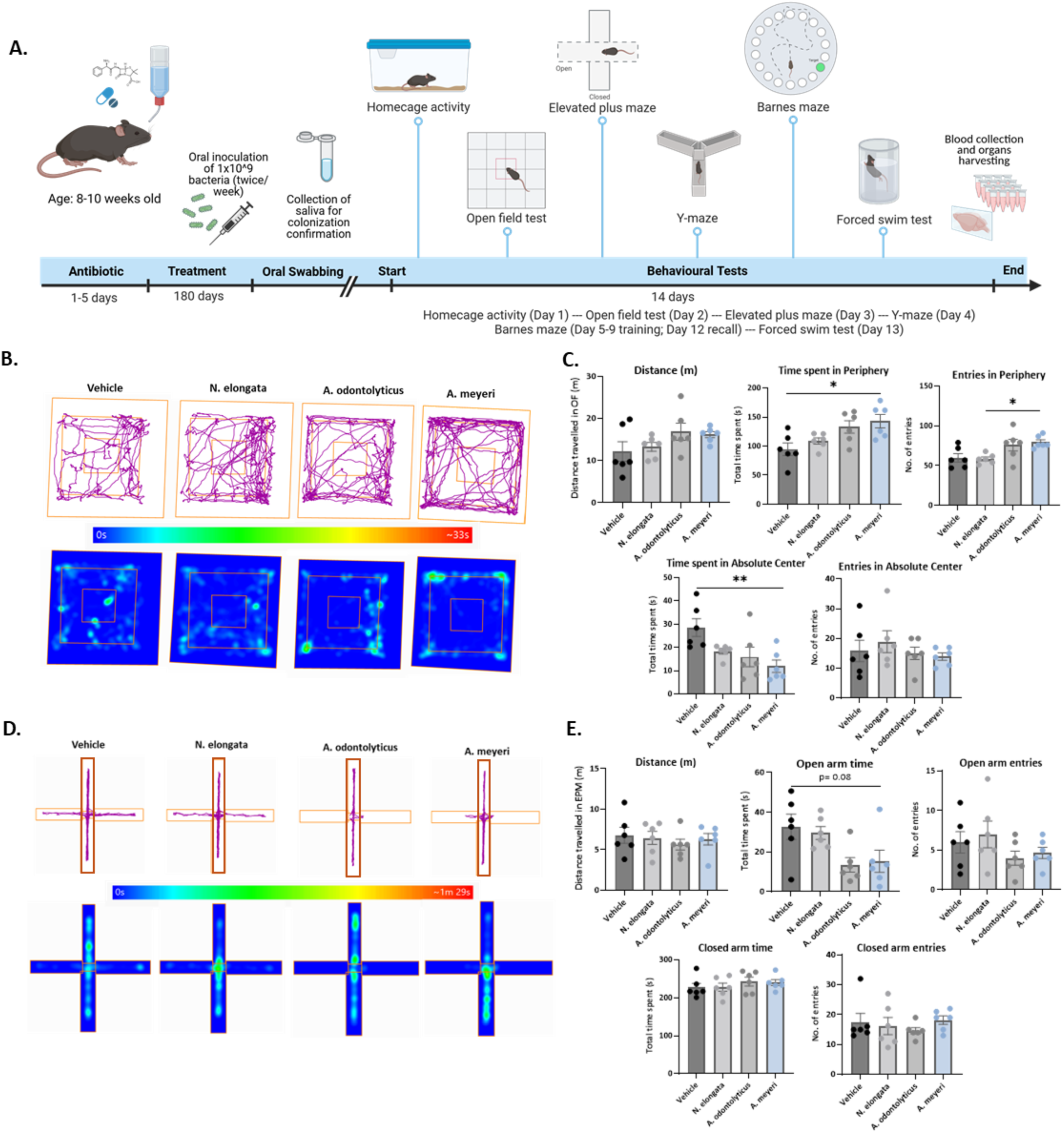
Chronic oral inoculation of *A. meyeri* induces anxiety-like behavior. **(A)** Schematic of the mouse study design. Following five days of oral antibiotic pretreatment, mice were chronically inoculated twice weekly for six months with *Actinomyces meyeri*, *Actinomyces odontolyticus*, *Neisseria elongata*, or vehicle control (1 × 10^9 CFU/mL). Oral swabs were collected to confirm colonization. Behavioral testing was performed in order of increasing stress intensity, followed by sacrifice and tissue collection. **(B)** Representative track plots and heatmaps from the open field test (OFT), illustrating movement patterns and time spent in different zones. **(C)** Quantification of OFT parameters, including time and number of entries into the periphery and absolute center, and total distance traveled. *A. meyeri*–inoculated mice exhibited increased time in the periphery (F*_(3,20)_* = 4.907, p = 0.0103) and reduced time in the center (F*_(3,20)_* = 4.94, p = 0.0100), consistent with anxiety-like behavior relative to controls. They also exhibited increased entries in the periphery as compared to *N. elongata* (F*_(3,20)=_* 4.709, p = 0.0121) **(D)** Representative trajectories and heatmaps from the elevated plus maze (EPM), showing time spent in open versus closed arms. **(E)** Quantification of EPM parameters, including time and entries in open and closed arms, and total locomotion. *A. meyeri and A. odontolyticus*–inoculated mice (F*_(3,20)_* = 4.040, p = 0.0213) showed a trend toward reduced open-arm exploration (p > 0.05) compared with vehicle controls, supporting OFT results. Data are presented as mean ± SEM (n = 6 per group). *p < 0.05, \*\**p < 0.01;* one-way ANOVA with Tukey’s multiple comparisons test.

### Oral *A. meyeri* activates microglia, which could be independent of neuroinflammation

We next examined the effects of chronic oral *A. meyeri* on CNS myeloid cell activation. It has been previously established that peripheral macrophages can infiltrate the CNS following injury, infection, or any event that compromises the blood-brain barrier (*25*). We aimed to investigate whether peripheral macrophages also infiltrate alongside any effect on microglial activation resulting from the chronic oral inoculation of *Actinomyces species*. First, using chromogenic RNA *in situ* hybridization (ISH), IBA1 expression was monitored in sagittal mouse brain FFPE sections; microglia were activated in the *A. meyeri*-treated group compared to the microbiome (*A. odontolyticus* and *N. elongata*) and vehicle controls (Fig. 2A-D). Furthermore, the IBA1 signal was not specific to any particular brain region following *A. meyeri* inoculation (Supplementary Fig. S3). Microglial activation was validated using a multiplex assay (shown in the green channel, Fig. 2C). Additionally, TMEM119 was included as a distinguishing marker of macrophages (negative) versus microglia (positive) (*26*). With careful scanning of the brain sections, no prominent cluster of TMEM119 was detected that was not co-localized with the IBA1 (Fig. 2A), i.e., all Iba1+ cells were TMEM119+ (Fig. 2B). It was noteworthy that microglia were also activated to a lesser extent (with no detectable peripheral macrophages) in the *A. odontolyticus* (same genus control bacteria which was also enriched in cannabis smokers (*15*)), (Fig. 2A and 2B). Due to differential translational regulation, RNA levels may not always correlate with protein levels. Thus, we performed immunohistochemistry to assess IBA1 protein expression, which further validated our findings that microglia were activated in a non-region-specific manner in the brains of *Actinomyces species*-treated mice (Fig. 2C and D).

**Fig. 2.**
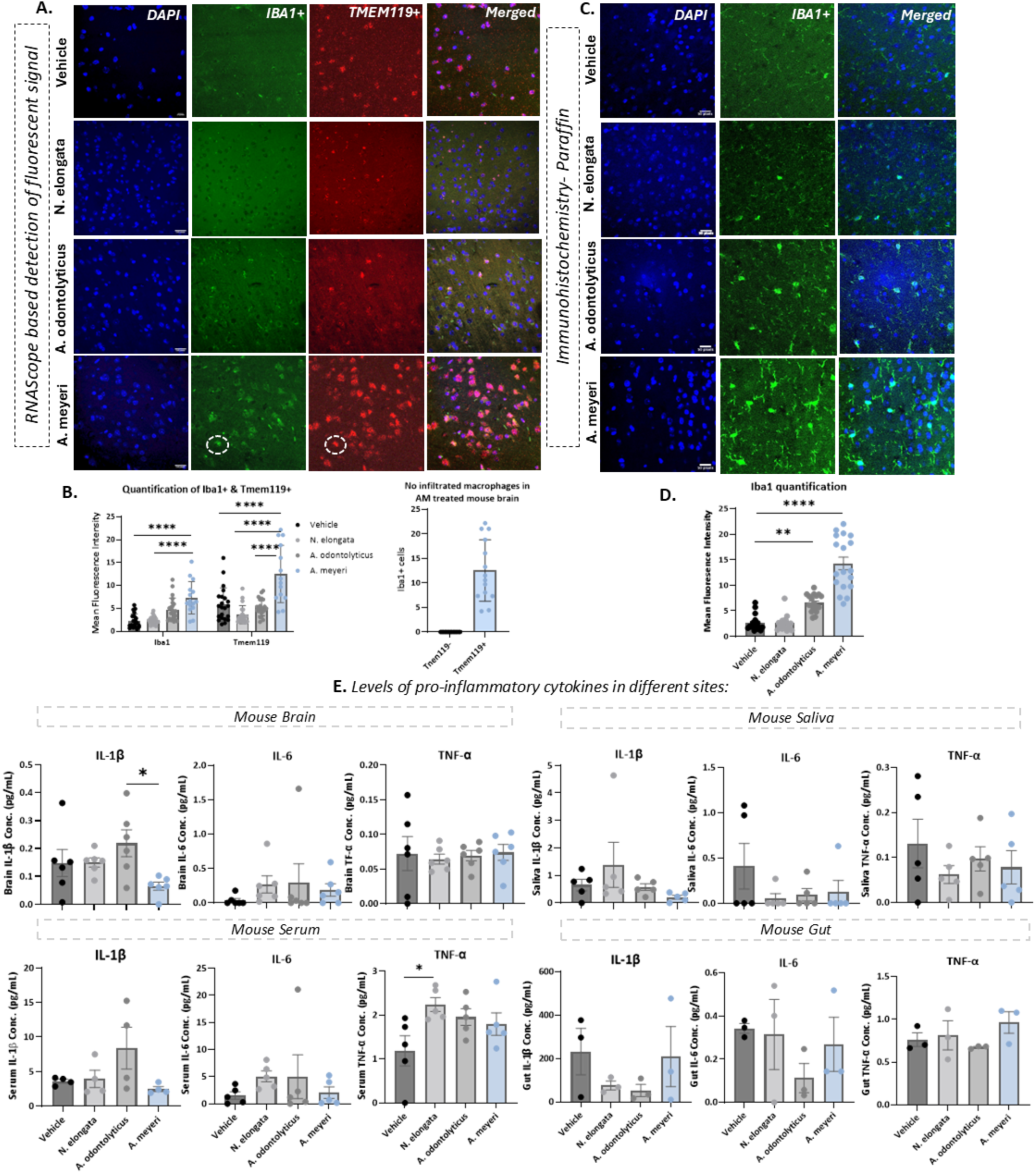
Chronic oral inoculation of *A. meyeri* triggers microglial activation independent of neuroinflammation and macrophage infiltration in the mouse brain. **(A)** RNA-based *in situ* hybridization (RNAScope). Representative images of 5 μm thin sections from the mouse sagittal brain embedded in FFPE. The scale bar is 100μm in the cortex of the brain. Sections were stained with IBA1 (activated microglia/macrophages, FITC) and TMEM119 (Red channel Cy3); a distinguishing marker for macrophages (negative) to microglia (positive), and DAPI as a counterstain. **(B)** Quantification of the mean fluorescent intensities (MFI) through ImageJ/Fiji from RNA-ISH studies. Microglia (Iba1 and TMEM119) activation was observed/increased in the *A. meyeri* inoculated group as compared to the control groups. Also, the MFIs were compared for IBA1 versus TMEM119. No prominent cluster of Iba1+ cells was detected that was not co-localized with TMEM119+, showing activation of microglia but not infiltration of systemic macrophages into the brain. **(C)** Immunohistochemical staining of IBA1. Representative images of 5 μm sections from a mouse sagittal brain embedded in FFPE. The scale bar is 50μm. Sections were immunostained with IBA1 antibody and DAPI as a counterstain. The Iba1 protein signal validated the activation of microglia at the protein level. **(D)** MFI analysis using ImageJ/FIJI of IBA1 immunohistochemical staining demonstrated increased microglial activation in the *A. meyeri*–treated group compared with vehicle controls. **(E)** Protein levels of the proinflammatory cytokines IL-1β, IL-6, and TNF-α were measured in the mouse brain, saliva, serum, and gut. No significant differences were observed between the *A. meyeri*–treated group and vehicle controls across these compartments, except for the IL-1β levels in the brain of the *A. meyeri*-treated group compared to the A. odontolyticus-treated group. Data are represented as mean ± SEM (n=3-6 in each group), *p < 0.05, **p <0.01, ***p <0.001, ****p <0.00001; One-way ANOVA, followed by Tukey’s multiple comparison test.

Neuroinflammation, an immune response in the brain, could be involved in various CNS-related disorders due to a variety of conditions (*27*). Interleukin-1β (IL-1β), Tumor necrosis factor (TNF-α), and Interleukin-6 (IL-6) are involved in various proinflammatory responses and diseases (*28–30*). While hypothesizing that chronic oral inoculation of *A. meyeri* could affect systemic as well as CNS inflammation, we collected serum, saliva, gut, and brain from mice at the end of the study. No significant changes indicative of inflammation were observed across the tissue types, i.e., brain, saliva, serum, and gut, as compared to the vehicle control (Fig. 2E). Unexpectedly, brain IL-1β levels following *A. meyeri* exposure showed a decrease compared to the *A. odontolyticus* group and a trending decrease versus *N. elongata* and vehicle controls (Fig. 2E), consistent with our scRNASeq data analyses of the brain samples (Figure S3).

### Oral *A. meyeri* inoculation results in CNS mitochondrial dysfunction and reduced GABA*ergic* neurotransmission

To study the transcriptomic landscape of various cell types in the brains of mice treated with *A. meyeri* (target oral bacteria), *A. odontolyticus*, *N. elongata* (bacterial controls), and vehicle control, the scRNA-Seq (10X genomics) technology was used (Fig. 3A). The single cells were isolated from the intact whole half-brains of individual mice from each study group. The estimated number of single cells obtained from each suspension was 14,000 to 20,000, whereas the target to capture was 3000-4000 nuclei in each integrated sample (Supplementary Fig. S4A). We identified 22 major cell types shown in different colors in *A. meyeri* vs Vehicle’s UMAP dimensional plots (Fig. 3B), while the other two UMAPs (*A. meyeri* vs. *A. odontolyticus* and *A. meyeri* vs *N. elongata*) and their subsequent downstream analysis are provided in Supplementary Fig. S4B-E. These includes, Somatostatin+ Interneurons (*Sst*), Layer 6 Intratelencephalic (IT) Neurons (*L6 IT*), Layer 6 Intratelencephalic Neurons (Car3) (*L6 IT Car3*), Layer 5 Extratelencephalic (ET) Neurons (*L5 ET*) LAMP5+ GABAergic Interneurons (*Lamp5*), Parvalbumin+ Interneurons (*Pvalb*), Layer 2/3 Intratelencephalic (IT) Neurons (*L2/3 IT*), Layer 6 Corticothalamic (CT) Neurons (*L6 CT*), Layer 5 Intratelencephalic (IT) Neurons (*L5 IT*), Astrocytes (*Astro*), Oligodendrocytes (*Oligo*), Meis2+ GABAergic Interneurons (*Meis2*), Oligodendrocyte Precursor Cells (*OPC*), Pericytes (*Peri*), Vascular-Associated Leptomeningeal Cells (*VLMC*), Endothelial Cells (*Endo*), Layer 5/6 Neocortical Projection Neurons (NP) (*L5/6 NP*), Layer 6b Neurons (*L6b*), Microglia and Paravascular Macrophages (PVM) (*Micro-PVM*), Synuclein Gamma (SNCG)+ Neurons (*Sncg*), Sst+ Chodl+ Interneurons (*Sst Chodl*), VIP+ Interneurons (*Vip*). Our results revealed that *A. meyeri* treatment led to a greater number of both up- and down-regulated genes in *L6IT, L6CT, L5ET, L5IT, L2/3IT* (glutamatergic excitatory pyramidal neurons), *Oligo* (glial cell; oligodendrocytes), and *Lamp5* (one of the several inhibitory GABAergic interneuron classes) compared to the other three study groups (Fig. 3B and 3C; Figure S4). Although we assessed microglial activation and cytokine levels in Fig. 2, Micro-PVM were not properly detected in the scRNASeq dataset due to technical limitations and thus were not specifically targeted here.

**Fig. 3.**
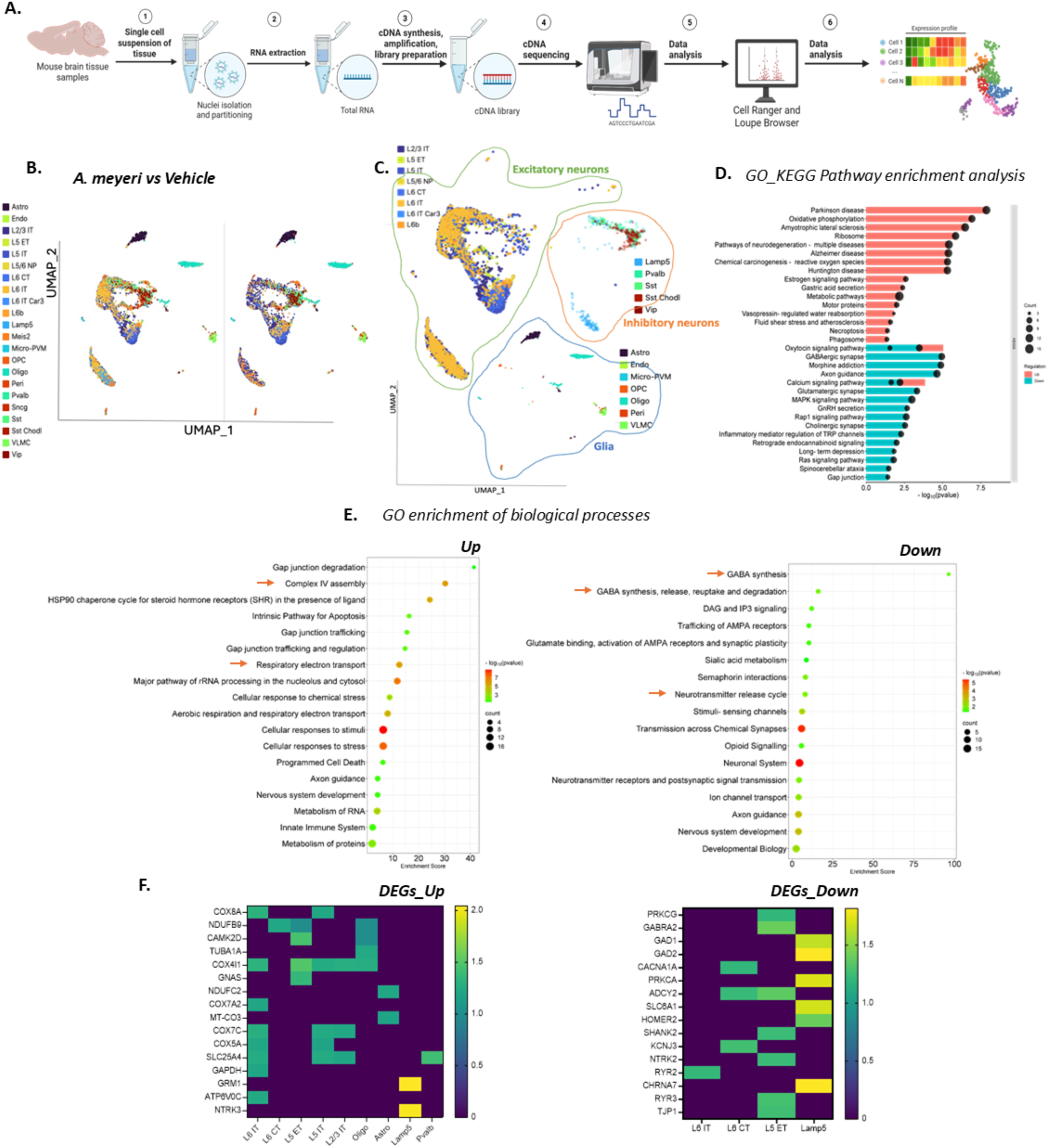
Chronic oral inoculation with *A. meyeri* induces broad transcriptional changes in neuronal cell types. **(A)** Schematic diagram illustrating the workflow for single-cell RNA sequencing (scRNA-seq) analysis of mouse brains. **(B)** Uniform Manifold Approximation and Projection (UMAP) dimensional plots showing the integrated dataset split by treatment with *A. meyeri* versus vehicle control. Twenty-two distinct cell types were identified by scRNASeq and are shown in different colors. **(C)** UMAP plot showing the classification of cell types into excitatory neurons, inhibitory neurons, and glial cells, along with their relative abundances and interactions. **(D)** KEGG pathway analysis of differentially expressed genes, highlighting upregulated (pink) and downregulated (blue) pathways, with circle size indicating gene counts and color scale representing –log10 *p*-values. **(E)** Gene ontology analysis of Reactome/biological processes using the DAVID annotation tool, displaying up- and downregulated pathways, enrichment scores [–log10(geometric mean of *p*-values)], gene counts (circle size), and –log10 *p*-values (color-coded). Pathways related to neurodegenerative disorders, oxidative phosphorylation/respiratory electron transport, and mitochondrial dysfunction were significantly upregulated. In contrast, pathways associated with GABAergic neurotransmission (synthesis, release, and reuptake) were selectively downregulated in *A. meyeri*–treated mice compared with controls. **(F)** Heatmaps representing the up-and downregulated differentially expressed genes (according to the average log_2_ fold-change) in different cell types in each integrated dataset. Neuronal cell populations exhibited a greater number of transcriptional alterations than non-neuronal cells.

Horizontal stack bar plot with dotline was prepared through the gene ontology KEGG pathway analysis, showing the overlap of up- and down-regulated genes in biological pathways as well as differently expressed (either up- or down) pathways (Fig. 3D); neurodegeneration relevant pathways were mainly up-regulated, while pathways linked to G-protein coupled receptors, GABAergic synapses were down-regulated in *A. meyeri* group versus control microbiomes and vehicle control (Fig. 3D and Supplementary Fig. S4C). Next, we looked into the gene ontology enrichment of the biological processes in the same integrated dataset (Enrichment bubbles in Fig. 3E). Enrichment score (ES) in the DAVID bioinformatics tool is a summary metric that helps prioritize relevant pathways based on their biological significance. The ES is a modified geometric mean (in log scale) of the p-value associated with the annotations within a given cluster or cell type. A higher enrichment score indicates that the cluster/cell type contains more statistically significant terms overall. For instance, an ES of 30 for complex IV assembly (part of the mitochondrial electron transport chain) in DAVID indicates robust statistical enrichment of that biological process in our differentially expressed gene list. This suggests that the input differentially expressed gene list contains many genes associated with Complex IV assembly, and it is highly unlikely to occur by chance. Many critical biological processes were captured in the transcriptomic landscape affected by the chronic oral treatment of *A. meyeri*. However, mitochondrial dysfunction (Fig. 3E; *Up-regulated*) and GABAergic neurotransmission (Fig. 3E; *down-regulated*) stood out with striking consistency (*A. meyeri*-treated group versus the other three study control groups) (Fig. 3E and Supplementary Fig. S4C). For instance, complex IV assembly, oxidative phosphorylation, and the respiratory electron transport chain are mostly flagged under the aspect of mitochondrial involvement. Similarly, the down-regulated gene ontology enrichment analysis of biological processes (BP) revealed a coordinated downregulation of multiple components of GABAergic neurotransmission following oral *A. meyeri* exposure. This included suppressed expression of genes involved in GABA biosynthesis, synaptic release, reuptake via transporters, and enzymatic degradation, suggesting a broad impairment in inhibitory signaling within the CNS. Overall, this dual pattern, i.e., enhanced mitochondrial activity coupled with suppressed GABAergic signaling, appears to be a specific and defining feature of the host response to oral *A. meyeri* treatment.

To further dissect cell-type-specific impact of *A. meyeri* treatment, we examined the distribution of differentially expressed genes (DEGs) across the 22 cell types. Heatmaps of up-and downregulated DEGs revealed that the majority of the transcriptional changes were concentrated in excitatory neuronal subtypes (*L5 ET* and *IT, L6 CT* and *IT, L2/3 IT*) and Lamp5-expressing GABAergic inhibitory interneurons, with additional but limited involvement of oligodendrocytes, astrocytes, Meis2+ GABAergic interneurons (Fig. 3F). Most of the mitochondrial markers or stress genes (e.g., COX genes, NDUF, SLC25 family) were upregulated in the excitatory neuronal subtypes in the *A. meyeri* treated group; some of them are up regulated in the other cell populations like oligodendrocytes and astrocytes.

GAD1 and GAD2 (glutamate decarboxylase 1 and 2) are rate-limiting enzymes for neuronal GABA synthesis. GAD1 is primarily involved in cytoplasmic GABA synthesis for metabolic purposes and extrasynaptic release, while GAD2 is more involved in regulating the vesicular pool of GABA for synaptic release (*31*). In GABAergic cells, SLC6A1 encodes the GABA transporter 1 (GAT-1), which is crucial for clearing GABA from the extracellular space (*32*). In the heatmap of DEGs_Down for *A. meyeri* compared to vehicle control (Fig. 3F), GAD1, GAD2, and SLC6A1 (or GAT-1) were downregulated in Lamp5, which is a GABAergic neuronal cell type. GABRA2 encodes the α2 subunit of the GABA-A receptor (postsynaptic), which is downregulated in Layer 5 ET (excitatory glutamatergic cell type). In our results, downregulation of GAD1/2 and SLC6A1 indicates reduced GABA ‘synthesis’ and ‘reuptake’, suggesting that these interneurons may be less active or exhibit diminished inhibitory function. Layer 5 ET neurons are glutamatergic, excitatory projection neurons that integrate inhibitory input from local interneurons (e.g., LAMP5, PV, SST cells). The downregulation of GABRA2 suggests reduced capacity to respond to GABAergic inhibition. Therefore, even if GABA is released, the ET neurons are less sensitive to it and may trigger behavioral deficits (anxiety-like behavior) in mice.

### Potential translocation of bacterial metabolites from oral to brain, but not the bacterial DNA, RNA, or peptidoglycan (PGN)

To further investigate the mechanism of *A. meyeri*-induced anxiety and neuropathogenesis, we assessed potential mediators, including bacterial DNA, RNA, cell wall/membrane product PGN, or metabolites, which may translocate from the oral cavity to the brain. First, no detectable *A. meyeri-*specific RNA signal was found in any region of the brain using RNAScope-based *A. meyeri*-specific probe in the sagittal mouse brain FFPE sections (Fig. 4A). We also tested specific probes for *A. odontolyticus* and *N. elongata*, and no signal was detected in the control microbiome-treated mouse brains (Supplementary Fig. S5A). Second, qPCR using specific *A. meyeri* primers was conducted, but no bacterial DNA was detected; purified bacterial culture DNA was used as a positive control (Fig. 4B). Bacterial-specific transgenes were included in the scRNASeq study; consistently, no transgenes of *A. meyeri* (or two other microbiomes) were found in the mice brains (Fig. 4C). After confirming the absence of bacterial DNA and RNA in brain, we evaluated bacterial PGN in the brain using the Fujifilm Wako chemicals kit. Bacterial PGN was undetected in the brain lysates, but was detected in the positive control using the lysed bacteria (Fig. 4D).

**Fig. 4.**
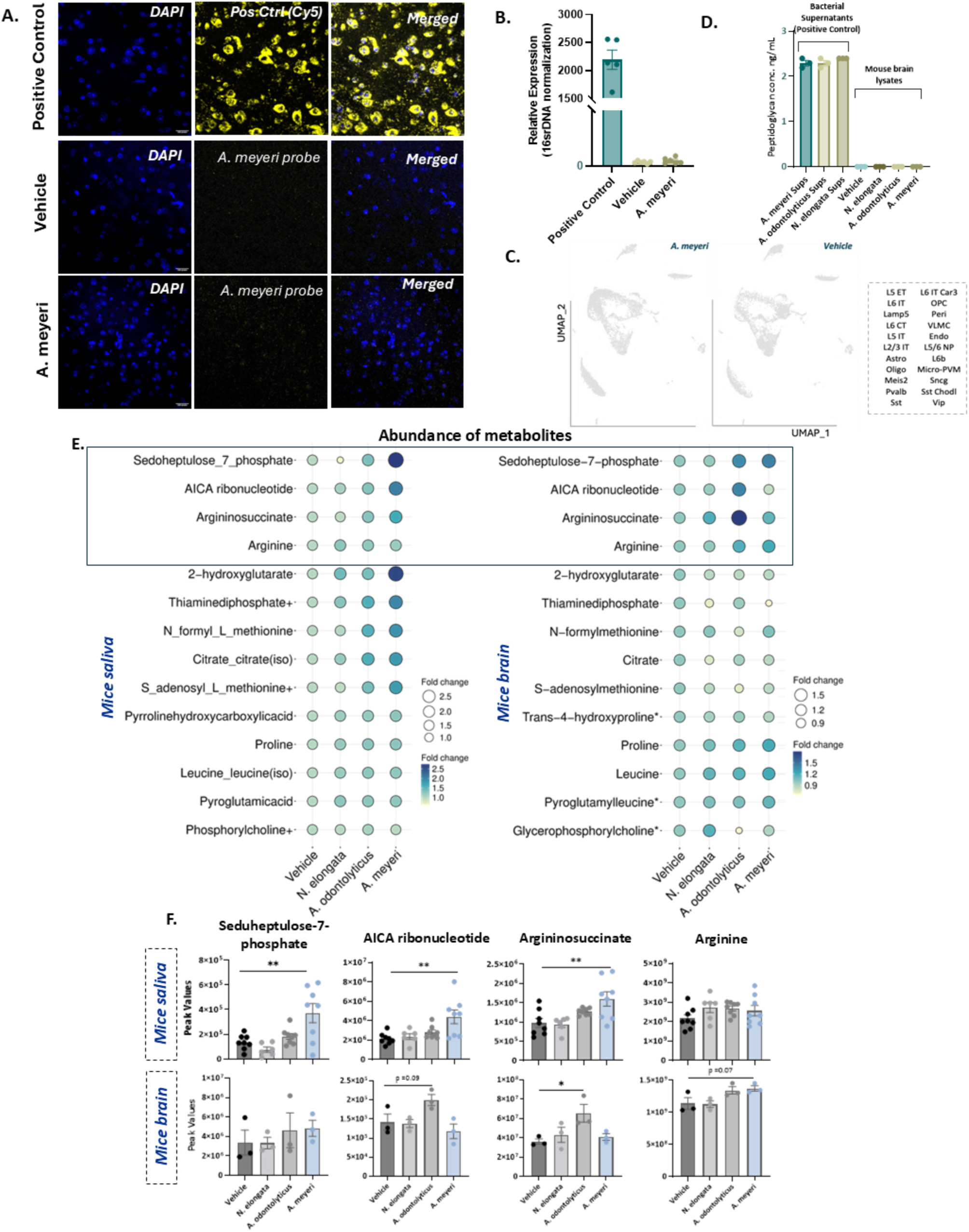
No oral bacterial entry into the brain, but metabolites are differentially abundant. **(A)** RNAScope in situ hybridization of sagittal mouse brain sections (5 μm FFPE). Fluorescent imaging showed no detectable *A. meyeri* RNA signal (yellow, Cy5 channel). A positive probe (UBC, a housekeeping gene) confirmed assay validity. Images were acquired at 40× magnification on a Zeiss LSM 510 Meta confocal microscope (1024 × 1024; n = 4). **(B)** Quantitative PCR analysis of brain lysates using *A. meyeri*–specific primers revealed no detectable bacterial DNA. Lysed bacterial cultures served as positive controls. **(C)** Targeted probe analysis in scRNA-seq (NCBI RefSeq: NR_029286.1) confirmed the absence of *A. meyeri-*specific RNA, further excluding bacterial translocation to the brain. **(D)** Peptidoglycan levels were determined in the brain lysates using the Silkworm larvae assay kit, with a pure bacterial culture serving as a positive control. **(E)** Metabolomic profiling illustrated by balloon plots. A total of 211 metabolites were detected in saliva and 919 in brain tissue. Twenty-three metabolites were significantly altered in saliva of *A. meyeri*–treated mice versus controls, 14 of which were also detected in brain tissue. Four metabolites were identified as candidate targets, showing parallel increases in saliva and brain in association with *Actinomyces* genus treatment. **(F)** Peak intensities of the four candidate metabolites are shown across saliva and brain in all three bacterial treatment groups. Data are presented as mean ± SEM (n = 3–8 per group). *p < 0.05, \**p < 0.01; one-way ANOVA with Tukey’s post hoc test*.

Microbiome dysbiosis occurs in many physiological conditions, like gut-related health issues, and has previously been associated with the altered levels of metabolites, for instance, short-chain fatty acids or indoles (*33*). It was hypothesized that microbes may communicate and exert influence on the host through the metabolites. To investigate this potential oral-to-brain mechanism, we conducted a metabolomics study to examine metabolic changes in both the oral cavity and the brain. A total of 211 metabolites were detected in the oral swabs, while around 919 metabolites (860 with known chemical identity and 59 unknown) were detected in the mouse brains (Supplementary Data: Figure 4). As we targeted the translocation of metabolites from the oral cavity, we found that 23 metabolites were significantly up-regulated in the oral swabs from *A. meyeri*-treated mice. Among those 23 metabolites, 14 were present in their paired brains as well (11 with exact chemical identity and 3 with slightly modified versions compared to the oral swab ones; shown in asterisks in the brain balloon plot). Although it would also be interesting to target the metabolites that were enriched in the oral swabs but reduced in their brains, we targeted the differentially increased metabolites in *Actinomyces* groups (Fig. 4E). We focused on identified four metabolites that were differentially abundant in the oral cavity and/or brain, i.e., Sedoheptulose-7-phosphate, 5-Aminoimidazole-4-carboxamide (AICA) ribonucleotide, Arginine, and Argininosuccinate (Fig. 4F). Sedoheptulose-7-phosphate, AICA ribonucleotide, and Argininosuccinate were significantly elevated in the oral cavity of *A. meyeri*-treated mice, whereas oral arginine levels remain unchanged. In contrast, within the brain, arginine was found to be trending increased in both Actinomyces microbiome treatments (also argininosuccinate in the *A. odontolyticus*), while other candidates did not show notable changes (Fig. 4F). The tissue-specific metabolite distribution highlights a potential shift in the arginine-argininosuccinate axis, suggesting compartmentalized regulation of this pathway. Furthermore, bacterial supernatants were screened to determine if these metabolites were being produced by the *Actinomyces* species bacteria or only by the host. Arginine/argininosuccinate was found in the *A. meyeri* and *A. odontolyticus* culture supernatants, confirming that this metabolite can also be produced by the two *Actinomyces* species bacteria (Supplementary Fig. S5B). Additionally, AICA ribonucleotide was increased in the brain of *A. odontolyticus* but not the brain of *A. meyeri*-treated group, and was not produced by any bacteria tested in vitro; Sedoheptulose-7-phosphate was increased in the oral cavity and the brain from both A. species-treated groups and was produced by *A. odontolyticus* but not *A. meyeri*, suggesting these two metabolites are not contributing to Am-induced behaviors or neuropathology. Based on this observation, in addition to AICA and Sedoheptulose-7-phosphate, we next focused on arginine and argininosuccinate as potential mechanisms of *A. meyeri*-induced mitochondrial bioenergetics and neurotransmitter balance, to determine whether their dysregulation may contribute to the observed neurobiological effects of *A. meyeri*.

### Hyperactive mitochondria affect the bioenergetics of the CNS cells with increased Reactive oxygen species (ROS) generation

We briefly mentioned in Fig. 3 that the upregulated pathways include oxidative phosphorylation, and the electron transport chain provided us with the list of genes that are upregulated. To visualize the mitochondrial alterations in the *A. meyeri*-treated group, we mapped the differentially expressed genes onto a schematic of the electron transport chain (ETC) (Fig. 5A). Genes encoding subunits of complex I (NDUFB9, NDUFC2) and complex IV/cytochrome c oxidase (COX4I1, COX5A, COX7A2, COX7C, COX8A) were significantly upregulated, along with MT-CO3, which is encoded by the mitochondrial genome rather than the nucleus. In addition, we observed upregulation of SLC25A4, an inner mitochondrial membrane carrier that mediates ADP/ATP exchange but does not directly belong to any respiratory complex (*34*). Although ATPV06, a subunit of the vacuolar ATPase, was also upregulated, it was not included in the schematic because it localizes primarily to lysosomal membranes rather than mitochondria (*35*). Together, these transcriptomic changes lead to a coordinated remodeling of mitochondrial respiratory chain and nucleotide transport components. From the transcriptomic screen, we selected COX4I1 and SLC25A4 for validation based on their functional relevance to mitochondrial activity. COX4I1 encodes a nuclear subunit of cytochrome c oxidase (Complex IV), the terminal electron acceptor in the respiratory chain, whereas SLC25A4 encodes the adenine nucleotide translocase (ANT1), which exchanges ADP and ATP across the inner mitochondrial membrane and is critical for bioenergetic homeostasis (*34*). The UMAP plots illustrate the upregulation of these genes mostly in the excitatory neurons (L2/3IT, L5, L6 subtypes) and oligodendrocytes (Fig. 5B). Western blotting confirmed a significant increase in SLC25A4 protein expression in the *A. meyeri*–treated group compared to vehicle controls, consistent with the transcriptomic findings; COX4I1 protein levels showed only a trend toward upregulation (Fig. 5C-D). These results support SLC25A4 as a robust marker of mitochondrial remodeling in response to *A. meyeri*.

**Fig. 5.**
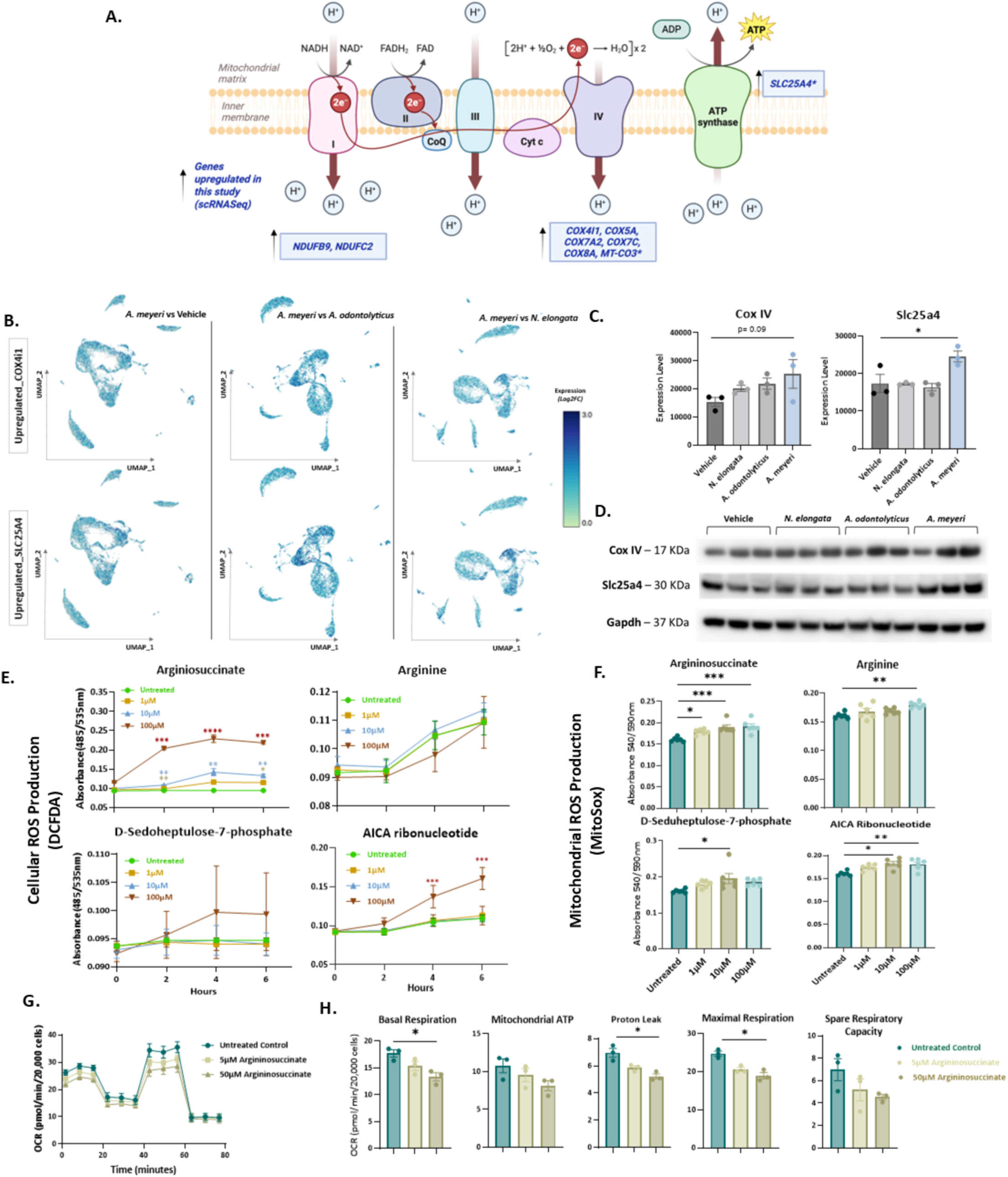
Chronic oral inoculation with *A. meyeri* impairs mitochondrial function, revealed through transcriptomic, proteomic, and metabolite-based functional assays. **(A)** Schematic highlighting upregulated genes in mitochondrial respiratory complexes I and IV (with *MT-CO3* encoded by mitochondrial DNA, others by nuclear DNA) and SLC25A4 (inner membrane ADP/ATP carrier). These changes suggest a compensatory response to energy deficit or stress, shifting mitochondria toward apoptosis. **(B)** UMAP plots from scRNAseq (using *A. meyeri* as reference) showing two scRNASeq shortlisted genes, i.e., COX4I1, a nuclear encoded subunit of complex IV (cytochrome c oxidase) in the electron transport chain (ETC), and SLC25A4 (or ANT1), a ADP/ATP translocator in the inner mitochondrial membrane (not part of ETC). These genes are shown to be upregulated in the excitatory neurons (L2/3IT, L5, L6 subtypes) and oligodendrocytes. **(C–D)** Western blot validation revealed a trend toward increased Cox IV expression and a significant increase in Slc25a4 protein, consistent with scRNAseq findings. **(E–F)** Cellular ROS (DCFDA/H2DCFDA) and mitochondrial ROS (MitoSOX) assays in N2a cells treated with metabolites (arginine, argininosuccinate, sedoheptulose-7-phosphate, AICA ribonucleotide; 1, 10, 100 μM). Argininosuccinate induced dose-dependent ROS generation at both cellular and mitochondrial levels. **(G)** Oxygen consumption rate (OCR) was measured in N2a cells treated with two concentrations of argininosuccinate under mitochondrial stress conditions (oligomycin, FCCP, and rotenone/antimycin A). **(H)** OCR analyses revealed effects on basal and maximal respiration, mitochondrial ATP production, proton leak, and spare respiratory capacity. Results were confirmed with technical triplicates. Data are mean ± SEM (n = 3–6 per group). *p < 0.05, **p < 0.01, ***p < 0.001, \*\*\**p < 0.00001; one-way ANOVA or mixed-model two-way ANOVA with Tukey’s post hoc test*.

Because our transcriptomic and protein analyses revealed remodeling of mitochondrial machinery (Fig. 5A-D), we next examined whether the shortlisted metabolites altered by *Actinomyces species* exposure (Fig. 4E and F) could themselves perturb mitochondrial homeostasis. We treated Neuro2a cells with various concentrations (1, 10, and 100 μM) of metabolite and assessed their effects on cellular and mitochondrial reactive oxygen species (ROS) production, as well as mitochondrial respiration. Using the DCFDA assay (Fig. 5E), argininosuccinate elicited a dose-dependent increase in total cellular ROS, whereas AICA ribonucleotide induced an increase only at the highest concentration (100 μM) in a time-dependent manner. MitoSOX analysis (Fig. 5F) further confirmed that argininosuccinate produced a robust, dose-dependent elevation in mitochondrial ROS, with additional effects observed for arginine (100 μM) and AICA ribonucleotide (10 and 100 μM). In contrast, sedoheptulose-7-phosphate did not affect total ROS but caused a modest increase in mitochondrial ROS at 10 μM. Given its consistent effects across both ROS assays, we next examined argininosuccinate in a mitochondrial stress test using the Seahorse XF analyzer (Fig. 5G-H). Treatment with argininosuccinate (5 and 50 μM) led to a dose-dependent suppression of mitochondrial respiration, with significant reductions in basal respiration, proton leak, and maximal respiration at 50 μM compared to untreated controls. Together, these findings highlight argininosuccinate, the most consistent microbiome-producing metabolite, which drives oxidative stress and mitochondrial dysfunction in neuronal-like cells, supporting its selection for further mechanistic investigation.

### Chronic oral *A. meyeri* exposure reduces GABAergic neurotransmission and lowers overall GABA content in the brain

Chronic oral inoculation with *A. meyeri* not only altered mitochondrial function (Fig. 5) but also exhibited anxiety-like behavior (Fig. 1). Given that mitochondria provide intermediates and essential metabolic precursors for neurotransmitter synthesis, including glutamate and GABA (*36, 37*), it is possible that mitochondrial dysregulation could impair GABAergic neurotransmission. As shown in Fig. 3, scRNASeq analysis indicated that the *A. meyeri* group exhibited reduced GABAergic signaling. Moreover, transcriptomic analysis (Fig. 6A) revealed downregulation of genes involved in multiple steps of GABA signaling: GAD1 and GAD2, SLC6A1 (GAT1), CACNA1A (involved in presynaptic release) and postsynaptic components GABRA2 and PRKCG/PRKCA (*38*). The UMAP plots demonstrated that GAD1, GAD2, and SLC6A1 were downregulated specifically in GABAergic neurons (Lamp5-positive) and astrocytes, with the *A. meyeri*-treated mice showing the most significant effect (Fig. 6C). Consistent with these transcriptomic changes, brain GABA levels (Fig. 6B) were significantly reduced in the *A. meyeri* group relative to other microbiome groups. Western blotting (Fig. 6D and E) confirmed decreased protein expression in vivo of GAD1 and GAD2, with GAT1 showing a trend toward reduction, reinforcing impaired GABA synthesis, reuptake, and signaling. A similar effect was seen in *A. odontolyticus* as well.

**Fig. 6.**
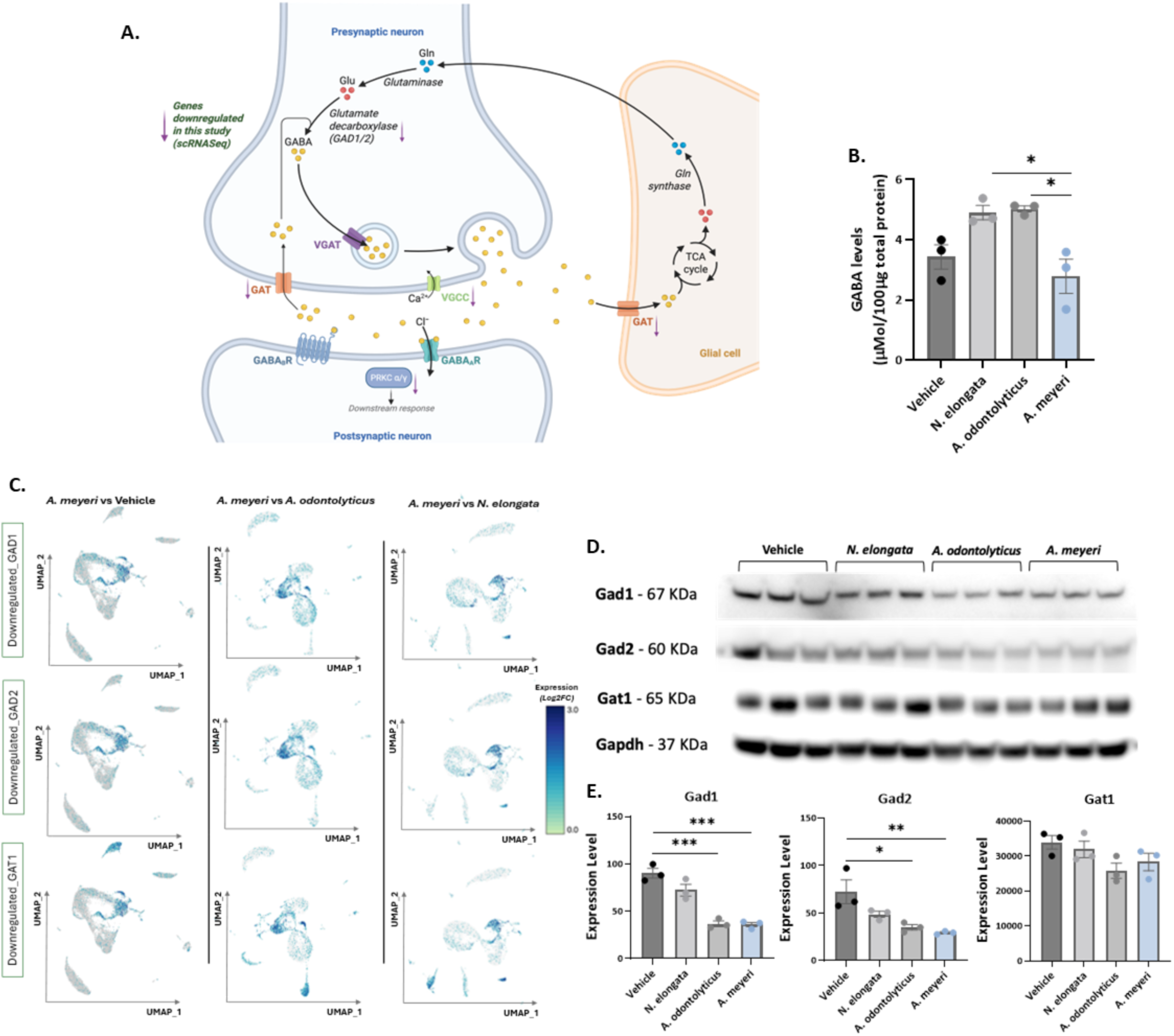
Chronic oral inoculation with *A. meyeri* reduces GABAergic neurotransmission. **(A)** Schematic illustrating the downregulation of genes associated with GABAergic neurotransmission. Reduced GABA synthesis was indicated by GAD1/GAD2 downregulation; impaired reuptake and recycling by reduced SLC6A1 (GAT1); diminished presynaptic release by CACNA1A downregulation; and weakened postsynaptic signaling by decreased expression of GABRA2, PRKCG, and PRKCA. **(B)** Quantification of GABA levels in mouse brain homogenates, after normalizing with the total protein content in each sample, revealed significant reductions in *A. meyeri*–treated mice compared with other microbiome groups. **(C)** UMAP plots (between-group comparison with *A. meyeri* group as reference) identified three downregulated genes in GABAergic neurons (Lamp5) and astrocytes: GAD1 (glutamate decarboxylase 67), GAD2 (glutamate decarboxylase 65), and SLC6A1 (GAT1). **(D–E)** Western blot validation demonstrated decreased GAD1 and GAD2 protein levels, with a trend toward reduced GAT1 expression in both *Actinomyces* groups relative to vehicle controls. Data are presented as mean ± SEM (n = 3–6 per group). *p < 0.05, **p < 0.01, ***p < 0.001, \*\*\**p < 0.00001; one-way ANOVA with Tukey’s post hoc test*.

Although Western blotting revealed that *A. odontolyticus* also reduced GAD1 and GAD2 protein levels and showed a trending decrease in GAT1, the scRNASeq analysis detected significant downregulation of GABAergic genes exclusively in the *A. meyeri* group (Fig. 6A and C). Consistent with these molecular findings, behavioral assays showed anxiety-like behavior only in *A. meyeri*-treated mice, whereas *A. odontolyticus*-treated mice exhibited a non-significant trend toward similar behavior (Fig. 1). Together, these results indicate that although the transcriptional, neurochemical, and behavioral alterations are more pronounced in *A. meyeri*, *A. odontolyticus* may have modest effects, highlighting the unique impact of *Actinomyces* species on inhibitory neurotransmission in the brain.

## DISCUSSION

Cannabis smoking is a widely consumed drug with a risk of various mental illnesses (*24*); however, the exact mechanisms are not fully understood. Previously, we found that the *Actinomyces* bacteria were highly enriched in the saliva of chronic cannabis smokers compared to non-smokers (*15*). We extended our investigations into whether the oral administration of cannabis-enriched bacteria in wild-type B6 mice could elicit behaviors indicative of potential neuropsychiatric illnesses in the absence of cannabis or its psychoactive components. In the current study, oral inoculation of *A. meyeri* in B6 mice for six months resulted in anxiety-like behavior, and activation of microglial cells without any prominent change in the canonical proinflammatory cytokines, i.e., IL-1 β, IL-6, and TNF-α. Moreover, the brains of *A. meyeri*-treated mice revealed hyperactive mitochondria and reduced GABAergic neurotransmission. We found no CNS bacterial or PGN translocation; however, components of the urea cycle and arginine/argininosuccinate-related metabolic changes were observed in the brain that induced mitochondrial dysfunction in Neuro2a cell line.

### Oral *A. meyeri* exposure induced anxiety-like behavioral deficits and microglial activation in wild-type mice

Oral microbiome not only plays a role in oral health but also can affect brain functions through the trigeminal nerve or olfactory system connections, as well as indirectly through the oral-gut-brain axis (*39*). The gastrointestinal tract contains a variety of bacteria whose metabolic products impact several physiological processes, such as stress, anxiety, depression, bowel diseases, and others. (*40*). In contrast, the role of the oral microbiome and its impact on the neurophysiological functions has only recently begun to emerge (*41*). The *Actinomyces* genus, preferentially an anaerobic or facultative-anaerobic bacterium, is not abundant in the gastrointestinal tract as compared to the oral cavity (*16*). In our previous investigations, we consistently failed to detect *A. meyeri* in the gut of mice orally inoculated with this bacterium for six months (*15*). This may be attributed to unfavorable gut conditions, including hypoxic gradients, limited nutrient availability, and competition with established microbial communities (*42*). Therefore, our results suggest that *A. meyeri* induced anxiety-like behavior via oral enrichment of *A. meyeri* and its metabolite translocation from the oral cavity to the brain, rather than the gut-to-brain microbiome.

Microglial activation and its related neuropathology can be independent of neuroinflammation, as microglia can adopt diverse functional states in response to various stimuli (*43*). These states range from classical proinflammatory responses (M1-like) to anti-inflammatory or neuroprotective phenotypes (M2-like), as well as metabolic and synaptic regulatory modes that can modulate neuronal function without eliciting overt cytokine signaling (*44, 45*). Consistent with this, RNAScope and IHC-P analyses revealed increased IBA1 expression in the two *A. species* groups, indicating microglial activation, while co-localization with TMEM119 confirmed these cells were resident microglia rather than infiltrating macrophages. Furthermore, levels of canonical proinflammatory cytokines (IL-1β, IL-6, and TNF-α) remained unchanged in brain, serum, saliva, and gut, indicating that the observed microglial response was restricted to the CNS and occurred independently of systemic and CNS inflammation. Together, these findings suggest that oral dysbiosis with *A. meyeri* triggers a localized microglial response that may contribute to neuronal and behavioral alterations without invoking classical inflammatory pathways.

### Oral *A. meyeri-*driven disruption of CNS mitochondrial function and GABAergic signaling

Our single-cell transcriptomic analysis revealed that chronic oral exposure to *A. meyeri* produces a dual signature of enhanced mitochondrial activity and suppressed GABAergic neurotransmission across discrete neuronal subtypes. Mitochondrial hyperactivity is a newly recognized factor in neurodegenerative diseases like Parkinson’s (*46*). Rather than a late-stage decline, overactive mitochondria may trigger early neuronal damage by increasing oxidative stress and disrupting energy balance, fueling a cycle of degeneration (*46*). Upregulation of mitochondrial genes, particularly within excitatory pyramidal neurons in our study, suggests a state of hypermetabolic stress that has been implicated in oxidative damage (can generate excess ROS) and neuronal vulnerability in CNS disorders (*47, 48*). It has also been recently reported that increased expression of SLC25A18 (a mitochondrial glutamate carrier) is associated with Alzheimer’s disease (AD) by screening differentially expressed genes in the brains of AD patients and by validation in animal models of AD and Neuro2A cells (*49*). In parallel, downregulation of GAD1, GAD2, and SLC6A1 in Lamp5 interneurons, along with reduced GABRA2 expression in Layer 5 excitatory neurons, indicates impaired GABA synthesis, reuptake, and postsynaptic responsiveness. These coordinated alterations indicate a loss of inhibitory tone, ultimately leading to an excitatory-inhibitory imbalance. Such an imbalance is a well-recognized mechanism underlying anxiety, epilepsy, and autism spectrum disorders (*50–53*). Importantly, these transcriptomic alterations align with the anxiety-like phenotype observed in our behavioral assays, providing a mechanistic link between oral dysbiosis, mitochondrial dysfunction, and disrupted neurotransmission.

Interestingly, comparison with bacterial controls revealed a genus-specific pattern. *A. meyeri* and *A. odontolyticus* produce similar metabolites, including neuropathogenic ones, in the oral cavity and the brain. While *A. meyeri* produced the most robust transcriptomic alterations, *A. odontolyticus* frequently trended in the same direction, albeit with weaker effects. This was particularly evident when integrating scRNA-Seq with protein-level validation and western blot analyses demonstrating significant reductions in GAD1, GAD2, and SLC6A1, as well as increases in COX-IV (Cox4i1) and SLC25a4, in both *A. meyeri* and *A. odontolyticus*, whereas scRNA-Seq detected these changes most strongly in the *A. meyeri* group. These findings suggest that while the *Actinomyces* genus as a whole may perturb mitochondrial and GABAergic pathways via neuropathogenic metabolites, *A. meyeri* elicits a more pronounced and consistent response. In contrast, *N. elongata,* depleted in cannabis smokers from our previous study (*15*), was indistinguishable from vehicle-treated controls across all assays, underscoring the specificity of *Actinomyces*-driven effects on brain function.

### Microbial metabolite-mediated neuropathogenic effects occur in the absence of bacterial translocation

Despite the behavioral, cellular, and transcriptomic alterations observed following chronic oral exposure to *A. meyeri*, we did not detect any evidence of bacterial translocation from the oral cavity to the brain. No traces of bacterial PGN were detected in the brains of treated mice. There is an ongoing debate in the field regarding whether oral bacteria can directly translocate into the brain or whether their systemic effects are mediated primarily through any soluble factor. While some studies have previously reported evidence for oral or gut bacterial presence (*P. gingivalis, F. nucleatum*) in the brain tissue under conditions of barrier disruption (*54, 55*), others argue that bacterial metabolites and immune mediators can be translocated to the brain and are sufficient to induce neural alterations without direct microbial invasion (*56–58*). Although metabolite concentrations may be low physiologically, chronic cannabis exposure may produce cumulative CNS effects (i.e., anxiety), consistent with the gradual progression of anxiety in chronic but not acute cannabis use. Metabolites can enter the brain through drinking water and alter CNS functions (*59, 60*). Our results also support an oral-brain axis model (*61*) in which soluble products derived from oral dysbiosis with *A. meyeri* influence neural processes (observed metabolite-driven effects on mitochondrial function) without the need for direct bacterial translocation. Notably, *Actinomyces* species are enriched in the saliva of chronic cannabis smokers without any clinical infections in our previous human study. In our current mouse study, *Actinomyces* species induced anxiety-like behavior and neuropathogenesis via bacterial metabolites, not bacterial translocation. This is consistent with oral-to-brain, rather than gut-to-brain, effects mediated by metabolite translocation, likely reflecting the shorter anatomical distance between the oral cavity and the brain, and consistent with evidence that acute cannabis use may reduce anxiety, but chronic heavy cannabis use induces anxiety. Thus, it takes many years to accumulate neuropathogenic effects in the brain via oral microbiomes and metabolites in the absence of direct bacterial translocation, providing a long window for the appearance of cannabis induced anxiety-related neuropathology.

Numerous metabolites were significantly enriched in the oral cavity of *A. meyeri and A. odontolyticus*-treated mice compared to the controls, and fourteen were detected in brain. Arginine and Argininosuccinate stand out in the *A. meyeri* group as central regulators of mitochondrial and redox homeostasis. They are also detected in bacterial supernatant, which can account for remote modulation of host metabolism. Arginine metabolism fuels multiple interconnected pathways, including nitric oxide (NO) synthesis, polyamine production, and the urea cycle, all of which directly influence neuronal excitability and survival. Argininosuccinate, as an intermediate in the urea cycle and connects nitrogen disposal with fumarate generation for the TCA cycle. Dysregulation of this axis has been linked to impaired ATP production and accumulation of ROS (*62, 63*), and elevated levels of urea have been reported in human dementia patients with Lewy bodies (*64*). The altered metabolism of arginine (*65*) and the role of the urea cycle in the brains of patients with AD (*66*) have also been previously reported. Such perturbations, like imbalances in arginine utilization, shift metabolism toward excessive NO or superoxide production, promoting peroxynitrite formation and redox imbalance (*67*). Elevated ROS is well established to impair mitochondrial integrity and has been implicated in both microglial activation and interneuron vulnerability (*68*). Moreover, AICA ribonucleotide, not detected in any bacterial culture supernatants, was increased in the brain of *A. odontolyticus* but not in the brain of the *A. meyeri*-treated group. Sedoheptulose-7-phosphate, produced by *A. odontolyticus* but not *A. meyeri*, was increased in brain from both *Actinomyces* species-treated groups, but not significant. These two metabolites can induce mitochondrial dysfunction at the highest concentration tested on ROS generation in neuro2a cells, suggesting that they may contribute to *A. odontolyticus*-induced neuropathology via host- or *A. odontolyticus*-derived metabolite translocation to the brain upon oral *A. odontolyticus* exposure. Our findings, therefore, provide a mechanistic link between *Actinomyces*-derived metabolite changes (arginine-argininosuccinate axis) and the mitochondrial dysfunction observed in our molecular assays.

Taken together, these results highlight that arginine-argininosuccinate dysregulation could be a potential metabolic bottleneck through which opportunistic oral commensal pathogens can remotely impair brain function. By disrupting this pathway, oral *Actinomyces* may set off a cascade of mitochondrial stress and oxidative damage that culminates in impaired GABAergic neurotransmission and anxiety-like behaviors. For example, in Drosophila, hyperactive mitochondria can sequester GABA through the mitochondrial transporter Aralar, leading to decreased GABAergic signaling and social behavior deficits (*69*) that might be our way forward to further investigating the interaction mechanism.

In summary, we report that oral *Actinomyces,* a previously defined chronic cannabis smoking-enriched oral microbiome, induces anxiety-like behavior in mice via impairing mitochondria and GABA signaling. We found that *Actinomyces* did not enter the brain and that most effects of the altered microbiome were attributable to metabolites. Thus, oral microbial metabolite-driven effects represent a critical but underappreciated component of the oral-brain axis, bridging local dysbiosis with remote behavioral and molecular phenotypes. However, given the complex interplay of metabolic networks, further research is warranted to delineate the precise mechanism through which the oral axis influences brain health and its potential interactions with other metabolic pathways.

## 2. MATERIALS AND METHODS

### 2.1. Experimental Animals

C57BL/6J mice were obtained from the Jackson Laboratories and housed in a specified ABSL2 facility with a controlled environment (temperature ranges 68-72 °F and humidity 54-58%), 12/12 hours day-light cycle with access to food and water *ad libitum*. The animal study protocol (IACUC-2019-00858-1 and IBC-2023-01589) was approved by the institutional committee for animal care and use, and all experiments were conducted according to the guidelines of the National Institutes of Health (NIH) for the care and use of laboratory animals. Before the start of the experiments, mice were kept in the facility for 5 days for acclimatization. Afterward, antibiotics (Amoxicillin Vancomycin cocktail 1mg/mL; MUSC pharmacy distribution center) were given to them in drinking water for five consecutive days to wipe out the oral microbiome, followed by the oral inoculation of the bacteria of interest. Building on our earlier study in female mice (*15*), we randomly assigned twenty-four female mice (aged 8-12 weeks) into four groups. They were orally inoculated with 1 × 10^9^ colony-forming unit (CFU) of *Actinomyces meyeri* (ATCC VPI 10648), *Actinomyces odontolyticus (BEI Resource, F0309)*, and *Neisseria elongata (ATCC NCTC 10660)*, as well as PBS or 2% Carboxymethyl cellulose (CMC) as vehicle control, twice per week for six months. We did not employ germ-free mice, as their immune immaturity and compromised mucosal defenses markedly increase susceptibility to bacterial colonization in the gut, which would confound our goal of studying oral-restricted exposure. Using conventionally colonized mice, therefore, allowed us to test oral inoculation effects without altering the broader microbial ecosystem or host immune tone.

Throughout the experiment, body weight changes were monitored weekly, and animal behavior was conducted in a balanced design to cancel out the environmental changes, time, and order effect. Saliva samples were collected after six months via PBS-soaked cotton swabs to determine the colonization (copy number) of bacteria of interest in the oral cavity. At the end of the experiments, mice were anesthetized with 5% isoflurane, and targeted tissues and fluids were harvested carefully. Serum corticosterone levels (Abcam AB108821 corticosterone ELISA) were assessed to monitor their physiological response to sociopsychological stress during the experiments.

### 2.2. Behavioral tests

#### 2.2.1. Home cage locomotor activity

Home cage activity was monitored for an hour to determine whether the oral bacterial treatment affected exploratory behavior or caused altered locomotion compared to vehicle control. Mice were acclimated to the environment 30 minutes before testing. Home cages were placed in an infra-red beam chamber (San Diego Instruments, San Diego, California), and the number of beam breaks was counted as a measure of locomotion. The automated system from SD instruments and PAS764 software recorded the animal movement via an infrared sensor.

#### 2.2.2. Anxiety-like behavior

Open field test (OFT) and elevated plus maze (EPM) test were performed to determine the anxiety-like behavior in mice.

The OFT apparatus was divided into three zones i.e., periphery (outer square), complete center (inner square), and absolute center (innermost square). Each mouse was placed in the absolute center as a starting point, and their locomotor activity was monitored for 5 minutes under a 120 Lux light source. ANY-maze software (Stoelting, Chicago, IL) tracked animal position and locomotor scoring.

The EPM test has four arms, i.e., two open arms (without the walls), two closed arms (with the walls), and a small center arena for placing the animal as a starting point. Each mouse was given 5 minutes under 120 Lux light sources after entering from the center arena, and the time spent in the center, open, and closed arms was determined by video recording and ANYmaze software.

#### 2.2.3. Spatial learning and memory

The Y-maze test and Barnes maze test were performed to determine the spatial working memory, spatial learning, and long-term reference memory, respectively.

The Y-maze test has three arms in a Y-shape, and the spontaneous alternations were determined as a measure of spatial working memory. Mice were placed into the center of the arms, and 5 minutes were given to them for exploring the maze under a 30 Lux light source. All three arms have a different visual cue to make them distinct from each other. ANYmaze software with a video source was used to calculate spontaneous alternations (the percentage of successive choices where the animals do not repeat an arm entry).

The Barnes maze apparatus is an elevated circular platform with twenty holes. A small, dark escape box is placed under the platform at one of the holes, which is referred to as an escape chamber. Four visual cues are placed on each of the four sides of the Barnes maze with different shapes and colors. This test was conducted under a bright overhead light (at least 450 Lux). All mice underwent a habituation session followed by training trials for five consecutive days, during which they were placed at the center of the maze and allowed to explore until they found the escape hole. If a mouse failed to locate the escape hole within 5 minutes, it was gently guided to the escape chamber. After training trials, the retention test trial was conducted following a two-day rest period, similarly to training. The latency to find the escape hole, number of errors, and search strategy were recorded through a video camera, and ANYmaze software was used to assess these measures of spatial learning and memory.

#### 2.2.4. Depression-like behavior

The forced swim test (FST) was performed at the end of the behavioral testing schedule. Mice were placed in the inescapable transparent tank filled with water (controlled temperature between 23-25 °C), and their escape-related mobility behavior was measured for 6 minutes through video recording. ANYmaze software was used to calculate the immobility time.

### 2.3. Bacterial quantification in mouse saliva

After a week of the last oral inoculation of *A. meyeri*, *A. odontolyticus*, and *N. elongata*, the mice were anesthetized, and oral swab samples were carefully collected from their tongue and gums using PBS-dipped cotton swabs. Total microbial DNA was extracted from the oral samples using a QIAmp DNA Microbiome kit (Qiagen) according to the manufacturer’s protocol. qPCR was performed using a CFX96 Real-time system (Bio-Rad). The target sequences were amplified using 10ng of DNA in a 20 μL reaction volume containing Perfecta SYBR Green SuperMix and specific primers (Supplementary Table 1). Each sample was tested in triplicate. The thermal parameters used for the reaction are as follows: 95°C for 5 min, 40 cycles at 95°C for 15 sec, and 60°C for 1 min. To count colony-forming units and construct a standard curve for each bacterium, genomic DNA was isolated from four serial dilutions of pure culture through the same method (starting from 1 × 10^9^). This genomic DNA was then used in the qPCR assay, and the C_t_ values obtained from the qPCR were employed to generate a standard curve and determine the CFU/mL titer of respective bacteria in mouse oral samples.

### 2.4. Detection of proinflammatory cytokines

Serum, saliva, brain, and gut samples were obtained from the control and treatment groups at the end of the study. IL-1β, IL-6, and TNF-α concentrations were determined in brain and saliva samples using the U-plex biomarker multiplex assay (Mesoscale Diagnostics), and cytokine levels in serum and gut were measured using an enzyme-linked immunosorbent assay (ELISA; Abcam). The respective detection procedure follows the manufacturer’s instructions.

### 2.5. Brain sample embedding and sectioning

Mouse brains were sagittally cut into two halves, and one half was used to embed in formalin-fixed paraffin-embedded blocks using a standard protocol. Tissue sections were cut (5µm) and mounted on charged slides. RNA *in situ* hybridization and immunohistochemical staining were performed as below:

#### 2.5.1. RNA *in situ* hybridization

The RNAScope 2.5 High Definition (HD) chromogenic duplex and multiplex fluorescent assays (ACDBio) were performed to visualize target RNA in the brain sections of control and treatment mouse groups. FFPE slides were pre-treated (including target retrieval step) according to the manufacturer’s protocol and followed by subsequent multiple steps of probe hybridization and signal development through HRP channel binding. The experiment consisted of hybridizing multiple probes, i.e., Iba1 (Mm-1089911), Tmem119 (Mm-472901), and unique 16s rRNA sequence probes for *A. meyeri* (B-1315481), *A. odontolyticus* (B-1328481), and *N. elongata* (B-1328471), along with positive (321811) and negative (321831) RTU control probes set. TSA vivid fluorophores were used to label different channels in the fluorescent assay, i.e., TSA plus fluorescein (green) with Iba1, TSA plus cyanine 3 (orange) with Tmem119, and TSA plus cyanine 5 (far red) with the respective 16s rRNA bacterial probe in a 1:1500 dilution. DAPI was used as a counterstain for nuclei. For the chromogenic assay, to demonstrate whether any brain region is specific to microglial activation, the Iba1 probe (1089911-C1) was used, and HRP-enzyme-based detection was performed. Hematoxylin-I was used as a counterstain, and stained sections were carefully mounted on the slides for visualization. Confocal microscopy was done using LSM Meta 550 and Leica Stellaris 8 for multiplex assay, while Leica Thunder’s bright-field microscopy was used for the chromogenic assay.

#### 2.5.2. Immunohistochemistry

Brain sections underwent deparaffinization and rehydration in different concentrations of xylene and ethanol, as previously described (*15*). After the heat-induced antigen retrieval step, the sections were washed with TBST and blocked with 10% normal serum (1% BSA in TBS) for an hour. Different sections were stained with Rabbit monoclonal to Amyloid precursor protein (ab32136) and rabbit monoclonal to Iba1 (ab178846) in a 1:500 dilution overnight at 4°C. After washing, goat anti-Rabbit IgG H&L (Alexa Fluor 488) was used (ab150077) in a 1:1000 dilution as a secondary antibody for an hour at room temperature. After the final wash steps, Vectamount mounting medium with DAPI (Vector Laboratories) was used, and the slides were cover-slipped. The Zeiss LSM 510 meta confocal microscope was used to visualize the signal. ZEN Software was used for image acquisition, and ImageJ for quantification of the signal.

### 2.6. Single-cell RNA Sequencing

Mouse brain samples from control and treatment groups were snap frozen and submitted for single-cell RNA sequencing to Singulomics Corporation (Bronx, NY). The nuclei were isolated from the frozen mouse brain tissue and constructed 3’ single-cell gene expression libraries (NEXT-GEN v3.1) using the 10X Genomics Chromium-X system. The libraries were sequenced with ∼200 million PE150 reads per sample on Illumina NovaSeq X Plus. The sequencing reads were analyzed with the mouse reference genome (GRCm39-2024-A). These reads include the addition of specific transgenes for detecting *A. meyeri*, *A. odontolyticus*, and *N. elongata* using Cell Ranger v8.0. Introns were included in the analysis.

The analysis workflow initiates with data processing through Cell Ranger, followed by rigorous quality control and additional data refinement. For the sample analysis, the Seurat workflow was employed, which includes normalization using the sctransform v2 pipeline. Subsequently, Louvain clustering was performed, enabling the identification of distinct cell populations within the sample. To further validate the cell types, the Azimuth application and Seurat’s anchor-based label transfer method were employed (*70, 71*). Besides that, cell types were identified using a reference dataset obtained from a similar tissue, specifically the mouse motor cortex. This reference dataset was constructed using data and annotations provided by the Allen Brain Institute and encompasses 159,378 nuclei integrated across 12 individual mice (*72*).

After sample integration with the Harmony algorithm, we obtained the UMAP plots for each condition/treatment. Differential expression analysis was done across all conditions, and a statistical threshold was applied, i.e., adjusted p-value < 0.05 and log2 fold change >+/−0.25, and FDR *q-*value (Benjamini-Hochberg) ≤ 0.05 were defined as differentially expressed genes (DEGs). For pathway analysis, the Gene Set Enrichment Analysis (GSEA) method was employed to enhance our understanding of gene expression patterns. The Wilcoxon Rank Sum Test was used to compare gene expression across conditions using the Seurat FindMarkers() function. The DAVID bioinformatics tool was used for functional annotation of KEGG and Reactome pathways, including the visualization of DEGs on pathway maps.

### 2.7 Estimation of Brain GABA levels

GABA levels in mouse whole brain lysates were quantified using a commercially available ELISA kit (Biomatik, EKF58282). Protein concentrations were first determined using the BCA assay, and sample dilutions were optimized to ensure accurate detection within the assay’s dynamic range.

Following ELISA quantification, raw GABA concentrations (pg/mL) were adjusted according to the dilution factor and normalized to total protein content. Final GABA levels were expressed as micro-molar (µM) per 100 µg of total protein, allowing for direct comparison with existing literature (*73*).

### 2.8 Detection of peptidoglycan in the brain

Bacterial peptidoglycan in mouse brain lysates was determined using a commercially available Silkworm Larvae Plasma (SLP)-HS Single Reagent Set Ⅱ (FUJIFILM Wako Chemicals). The assay principle is that the hemolymph of the silkworm Bombyx mori contains a self-defense mechanism known as the “prophenoloxidase (proPO) activating system” or “proPO cascade”. Upon invasion by a microbe, the system participates in melanin formation, as observed in the body fluids of insects, to protect them from the invader’s attack. The system is activated by peptidoglycan (PGN) from bacteria, and consequently, proPO in the system is activated. The system is thought to be a cascade reaction involving the activation of multiple zymogens of proteinases. The SLP reagent is a lyophilized product containing factors of the proPO cascade. It is prepared from the body fluid of silkworm larvae and processed without melanin formation. PGN activates the SLP reagent, then L-3,4-dihydroxyphenylalanine (DOPA) is oxidized, and melanin is formed. Since PGN is found not only in gram-positive but also gram-negative bacteria, the SLP reagent responds to a wide range of bacteria, regardless of their Gram-staining classification. Therefore, this kit is used to detect the presence of PGN by measuring the melanin pigment formation of the SLP reagent as an indicator. As a positive control, pure cultures of lysed bacteria (*A. meyeri, A. odontolyticus, and N. elongata*) were used.

### 2.9. Western Blotting

Mice whole brain lysates were prepared using RIPA lysis buffer (ThermoFisher 89900) supplemented with a protease/phosphatase inhibitor cocktail. Protein estimation of the samples was performed using the BCA method. An equal amount of protein (25μg) was resolved by SDS-PAGE and transferred to PVDF membranes (Millipore). Membranes were blocked with 5% skim milk in TBS-T, incubated with primary antibodies overnight at 4 °C, and then incubated with HRP-conjugated secondary antibodies (Cell Signaling Technology) the next day. Bands were visualized using ECL (Thermo) and quantified with ImageJ. Primary antibodies are Cox IV mouse mAb # 1967 (4D11-B3-E8, CST), ANT1 polyclonal Ab (PAS-109394, Thermo), GAD1 (#5305S, CST), GAD2 (D5G2) XP Rabbit mAb (#5843S, CST), and GAT1 Rabbit mAb (373425, CST). Secondary antibodies are anti-rabbit (Cat #7074S, CST) and anti-mouse (Cat# 7076S, CST).

### 2.10. Screening of metabolites in the brain, saliva, and bacterial supernatants

Mice saliva samples were collected as described in detail previously (*15*). Brain samples were harvested, and homogenization of frozen tissues was performed under liquid nitrogen. These samples were sent to Metabolon Inc. (Durham, NC) for the metabolomic study. Samples were prepared using the automated MicroLab STAR® system. Proteins in the sample were precipitated (with the aid of a small molecule matrix) using methanol under vigorous shaking for 2 minutes, followed by centrifugation. The resulting extracts were subjected to different chromatographic and spectrometric pipelines as previously described by Ford and colleagues, 2020 (*74*). Briefly, the untargeted metabolomics analyses were performed using ultrahigh-performance liquid chromatography/tandem mass spectrometry (UPLC-MS/MS) and gas chromatography-mass spectrometry (GC-MS), following their standard procedures. Reference libraries were used to identify different metabolites with their raw peak values. Peaks were quantified using the area under the curve.

*Actinomyces meyeri* and *odontolyticus* were cultured in the Actinomyces-specific broth (HIMEDIA M233) for 48 hours. Bacterial cells were pelleted down, and supernatants were filtered out (0.22µm) for the untargeted metabolomics study executed by Creative Proteomics Inc. through UPLC-MS. Separation was performed by ACQUITY UPLC (Waters) combined with Q-Exactive MS (Thermo) and screened with ESI positive and negative modes, provided with both Hilic and C18 columns.

### 2.11. In vitro assays of mouse neuroblastoma cell lines

Neuro2a (ATCC CCL-131) cell line was maintained in the DMEM medium (Cat # 11965-092) with 10% FBS and 5% PSN antibiotic solution (Cat # 15640-055) and grown until the cells looked healthy and confluent in cell culture-treated plates (Cat # 240304; 100×15mm) at 37°C and 5% CO2 in a humid atmosphere.

#### 2.11.1 DCFDA-ROS assay

The cellular reactive oxygen species (ROS) levels were determined using the DCFDA/H2DCFDA ROS assay kit (Abcam ab113851). TBHP 200µM and H2O2 400µM were used as positive controls. Briefly, Neuro2a cells were plated (seeding density 25,000 cells per well) in an FBS-supplemented DMEM without phenol red media (Cat # 21063-029) onto a 96-well black plate (Cat # 3603 Corning Costar). After 24 hours, cells were washed with 1x wash buffer (provided in the kit) and incubated with 10µM DCFDA for 45 minutes at 37°C (in the dark). The DCFDA solution was removed carefully, and different treatments of interest (metabolites in various concentrations) in the same FBS-supplemented, without phenol red DMEM media were added to their respective wells. ROS production was determined immediately by measuring the formation of fluorescent dichloro fluorescein (DCF), using a spectrophotometer, at 458/535 excitation/emission wavelengths for 6 hours (every 2 hours). The value of fluorescence intensity at each time point is reported in triplicate.

#### 2.11.2 MitoSox ROS-assay

The mitochondrial superoxide (MitoSox; refers to the reactive oxygen species (ROS) generated specifically in mitochondria) levels were determined using the Mitochondrial Superoxide fluorometric Assay Kit (abcam ab219943). Antimycin A 50µM was used as a positive control. The procedure of seeding the N2a cells was essentially the same as in the DCFDA assay; however, a 5 × 10^5^ seeding density per well was used. After 24 hours, the old media was gently replaced with the fresh complete media (without phenol red), containing treatments (different metabolites) or antimycin A, and incubated for 30 minutes at 37 °C in the dark for superoxide production. Afterwards, 100uL/well diluted MitoROS staining solution was added, and further cells were incubated at 37°C for 30 minutes protected from light. After the incubation. endpoint measurement of fluorescent intensities was monitored at Ex/Em = 540/590 nm. Each treatment was tested in six replicate wells.

#### 2.11.3 Mitochondrial function assays

Oxygen consumption rates (OCRs) were measured using an Agilent Seahorse XFe96-Pro analyzer according to the Agilent Mito stress test protocol. Neuro2a cells were seeded onto the Seahorse 96-well plate at a density of 40,000 cells per well for 2 hours after treatment with different concentrations of the metabolite of interest and then incubated for 22 hours. One hour before the assay, plated cells were incubated at 37°C in the absence of carbon dioxide. Cells were sequentially treated with mitochondrial poisons, i.e., Oligomycin (1.5µM), Carbonyl cyanide-4 (trifluoromethoxy) phenylhydrazone (FCCP) (1.5µM), Rotenone, and Antimycin (0.5µM) while measuring the OCR through the Seahorse instrument. (a) Basal respiration was determined by subtracting the non-mitochondrial OCR (after rotenone and antimycin A treatment) from the basal OCR (before any mitochondrial poison treatment), (b) ATP synthesis by subtracting the OCR after oligomycin from the baseline OCR, (c) Proton Leak by subtracting non-mitochondrial OCR from the OCR after oligomycin, (d) Maximal respiration by subtracting non-mitochondrial OCR from the OCR after FCCP, (e) Spare respiratory capacity by subtracting basal respiration from the maximal respiration. Phase contrast microscopy was performed on the plate to assess the cell confluency and confirm the equal seeding density through CellProfiler software for normalization of OCRs.

### 2.12. Statistical analysis

Statistical analysis was performed using GraphPad Prism version 10, and data are presented as mean±S.E.M unless otherwise specified in the respective figure legend. Comparison between the groups was performed using two-tailed unpaired Student’s t-test, one-way analysis of variance (ANOVA), two-way ANOVA, mixed model, or repeated measure design with Tukey’s multiple comparison tests. The data from ScRNASeq and untargeted metabolomics studies were analyzed using their respective bioinformatics software. Significance was defined as p< 0.05. Image acquisition and quantification (ImageJ/Fiji) were conducted in a blinded manner, with identical acquisition settings and standardized analysis parameters applied across all groups to ensure unbiased measurements. Information on the sample size, number of replicates, and the statistical test used for each experiment is included in the figure legends.

## ACKNOWLEDGMENT

◦ Behavioral testing was supported by the CNDD Mouse Behavior Phenotyping Core (P20GM148302), a University Shared Research Facility at MUSC.
◦ Image facilities were supported in part by the Cell & Molecular Imaging Shared Resource, MUSC Cancer Center Support Grant (P30 CA138313), the SC COBRE in Digestive and Liver Diseases (P20 GM130457), the MUSC Digestive Disease Research Cores Center (P30 DK123704) and the Shared Instrumentation Grants S10 OD018113 and S10 OD028663.
◦ Immunohistology work was supported in part by the Translational Science Shared Resource, Hollings Cancer Center, Medical University of South Carolina (P30 CA138313).
◦ Seahorse XF assay work was supported in part by the Bioenergetics Profiling Core, Medical University of South Carolina, for Seahorse XF Pro energy metabolism analyses.
◦ SRplot: A free online platform for data visualization and graphing.

## AUTHOR CONTRIBUTIONS

Conceptualization: WJ, SF; Methodology: TS, RDP, AAN, HX; Investigation: TS, ZL, DJ, BB, ZY; Data analysis and Visualization: TS, ZW; Supervision: WJ, SF; Funding acquisition: WJ, SF; Writing—original draft: TS; Writing—review & editing: TS, WJ, SF, RDP, JFM, PWK, HX, ZY.

## DATA AND MATERIAL AVAILABILITY

All data are available in the main text or the supplementary materials.

---

**Supplementary Table 1:**
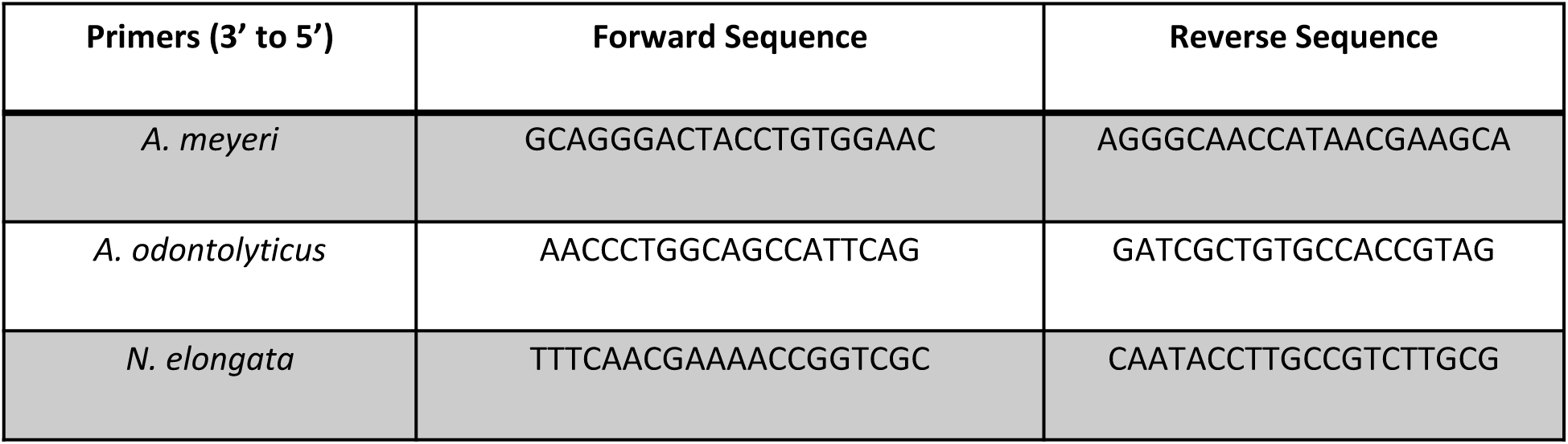
The oligos used for qPCRs.

**Supplementary Fig. S1:**
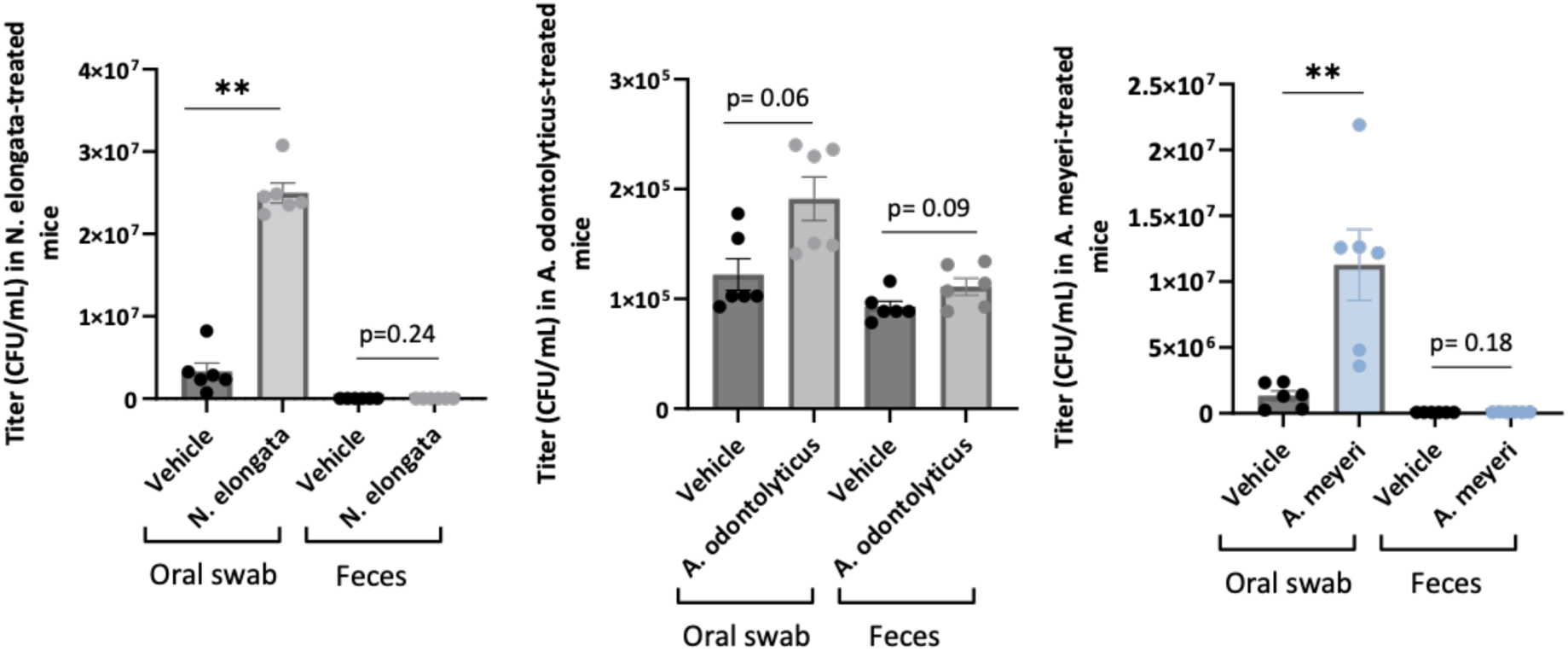
Quantification of bacterial DNA in saliva (oral swabs) and fecal (stool) samples after oral inoculations. The oral swab samples were collected from the mice after chronic oral inoculation of the microbiome, while feces were collected from the intestines at the time of mouse sacrifice (fresh catch). The quantification of *A. meyeri, A. odontolyticus*, and *N. elongata* specific DNA was evaluated using qPCR with their specific primers. To count colony-forming units and construct a standard curve for each bacterium, genomic DNA was isolated from four serial dilutions of pure culture through the same DNA extraction method (starting from 1 × 10^9^). This genomic DNA was then used in the qPCR assay, and the Ct values obtained from the qPCR were employed to generate a standard curve and determine the CFU/mL titer of respective bacteria in mouse oral samples. Data are represented as mean±SEM (n=6 in each group), **p < 0.001, One-way ANOVA, followed by Mann-Whitney U-test.

**Supplementary Fig. S2:**
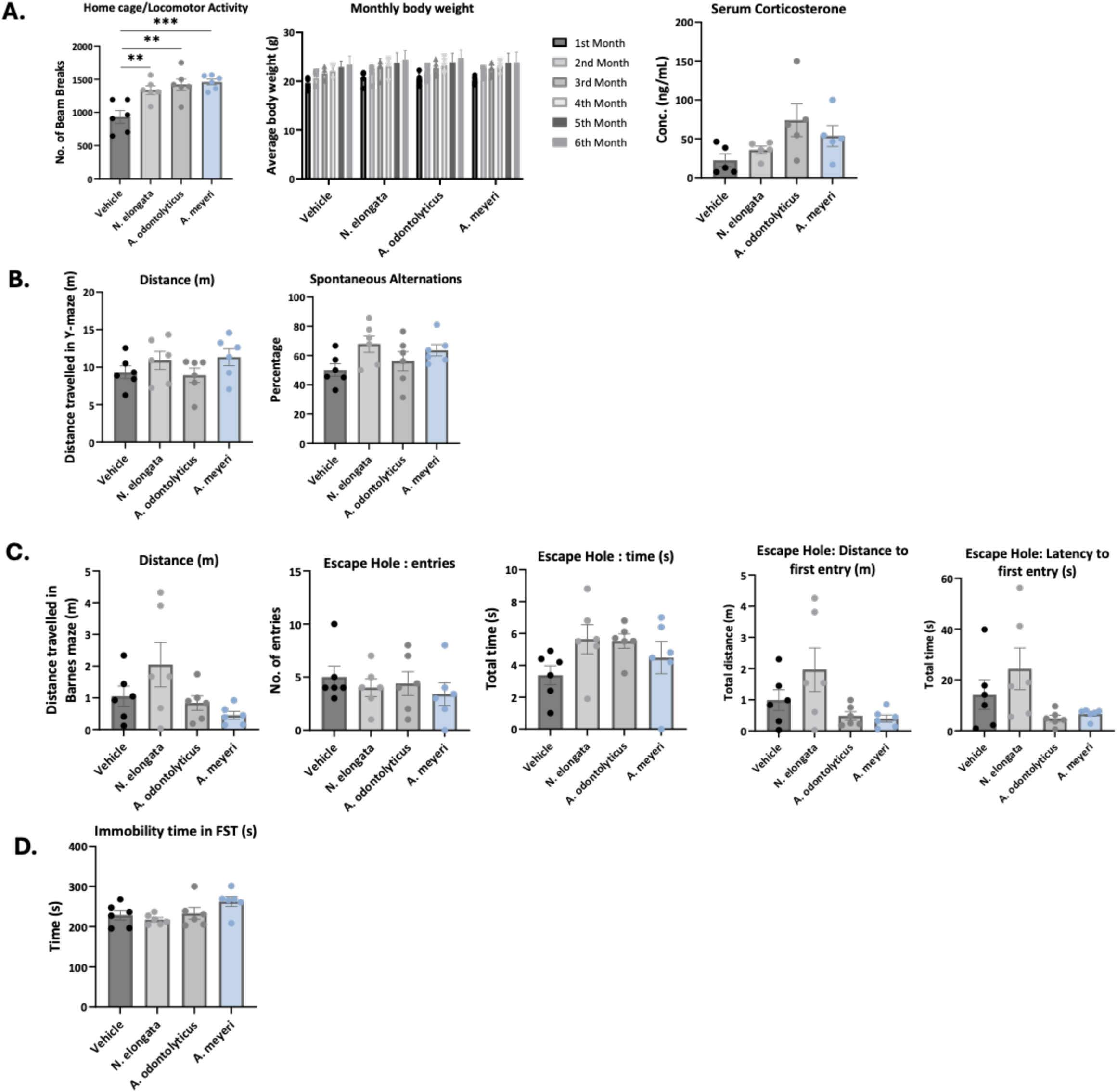
No prominent change in mouse behaviors related to cognitive functions, depression, and baseline serum corticosterone levels. **(a)** Baseline levels before starting the battery of behavioral tests. Home cage locomotor activity was increased in all microbiome treatments as compared to the control (F*_(3,20)_* =10.12, p=0.0003). All mice healthily gained their body weight during oral treatments of *A. meyeri, A. odontolyticus*, and *N. elongata*. The mice did not show any changes in their serum corticosterone levels, indicating no handling or environmental stress. **(b)** Y-maze activity showed no change in the spontaneous alternation in any of the microbiome groups as compared to the vehicle control. **(c)** Different parameters were tested in the Barnes maze on the recall day after extensive five-day training. No significant change was observed in any parameter in any of the microbiome groups. **(d)** The forced swim test showed a trending increase in immobility time in the *A. meyeri* treatment, but it was not significant. Data are represented as mean±SEM (n=6 in each group), **p < 0.01, ***p <0.001; One-way ANOVA, followed by Tukey’s multiple comparison test.

**Supplementary Fig. S3:**
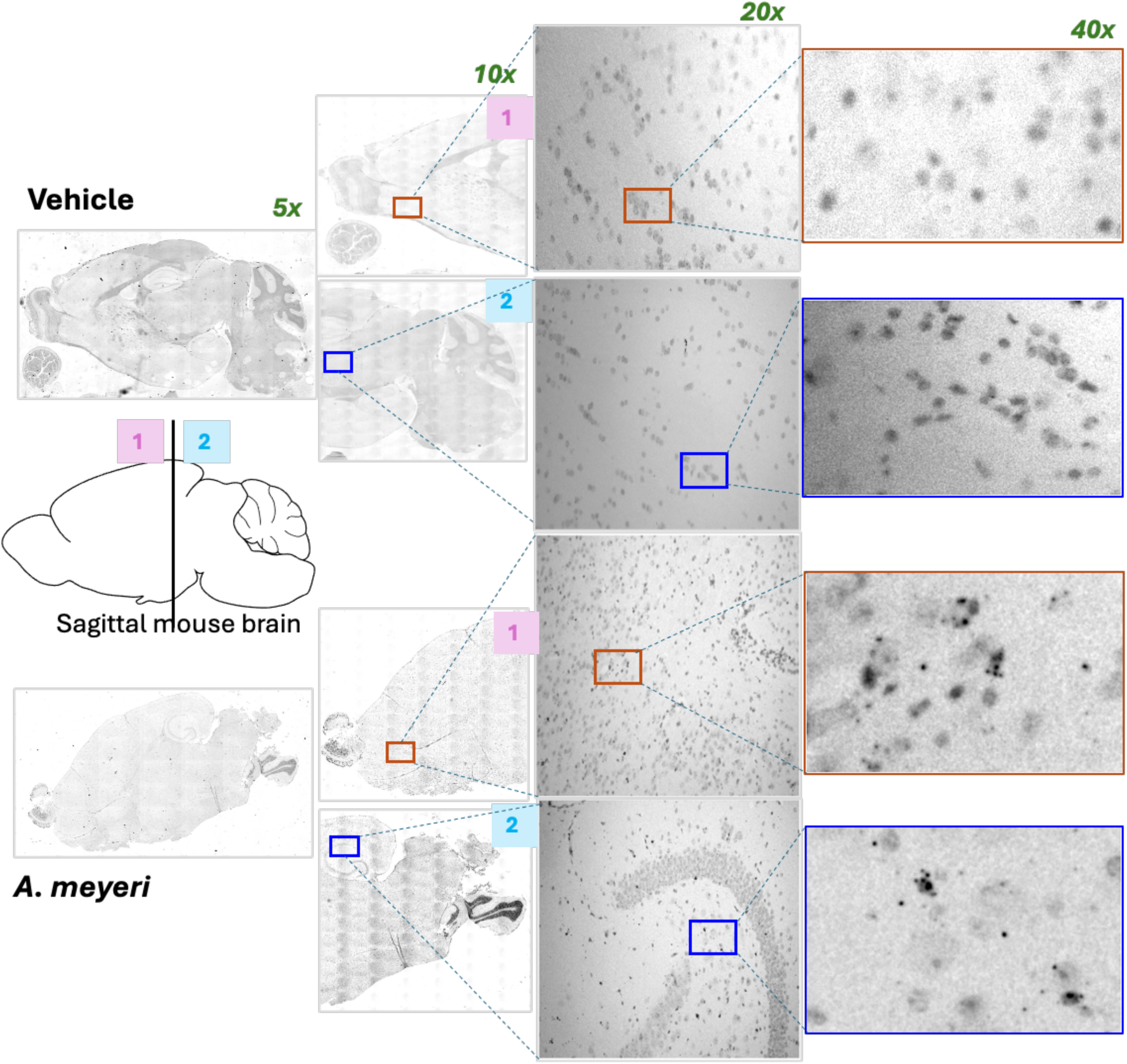
Non-region-specific activation of Iba1+ cells in the mouse brain after chronic administration of *A. meyeri*. ***RNA-based in situ hybridization:*** Representative images of mouse sagittal brain sections (5µm) that were treated with vehicle and *A. meyeri* for six months. These images were taken using a Leica Thunder (brightfield) microscope at 5x, 10x, 20x, and 40x magnification to illustrate the Iba1-positive signal (RNAScope probe # 1089911-C1) in various brain regions of *A. meyeri*-treated mice compared to the control group. The chromogenic signal was converted into grayscale through ImageJ/Fiji for better visualization. HRP enzyme-based detection was performed, and hematoxylin I was used as a counterstain. The random selection of the areas in the sagittal brain sections of the vehicle and *A. meyeri* treated groups is shown in the figure by zooming in from 5x to 40x. Samples were in triplicate in each group.

**Supplementary Fig. S4:**
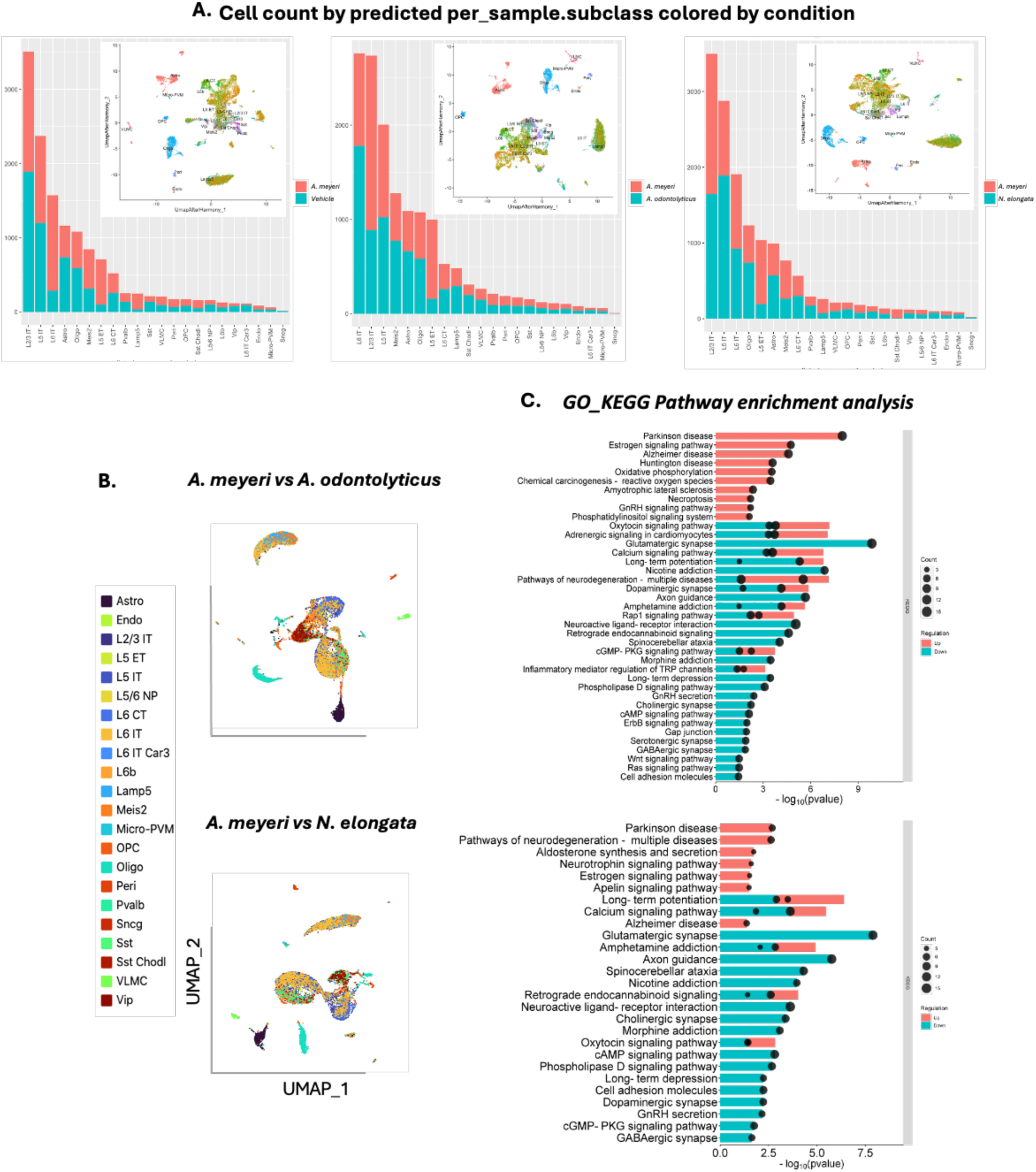
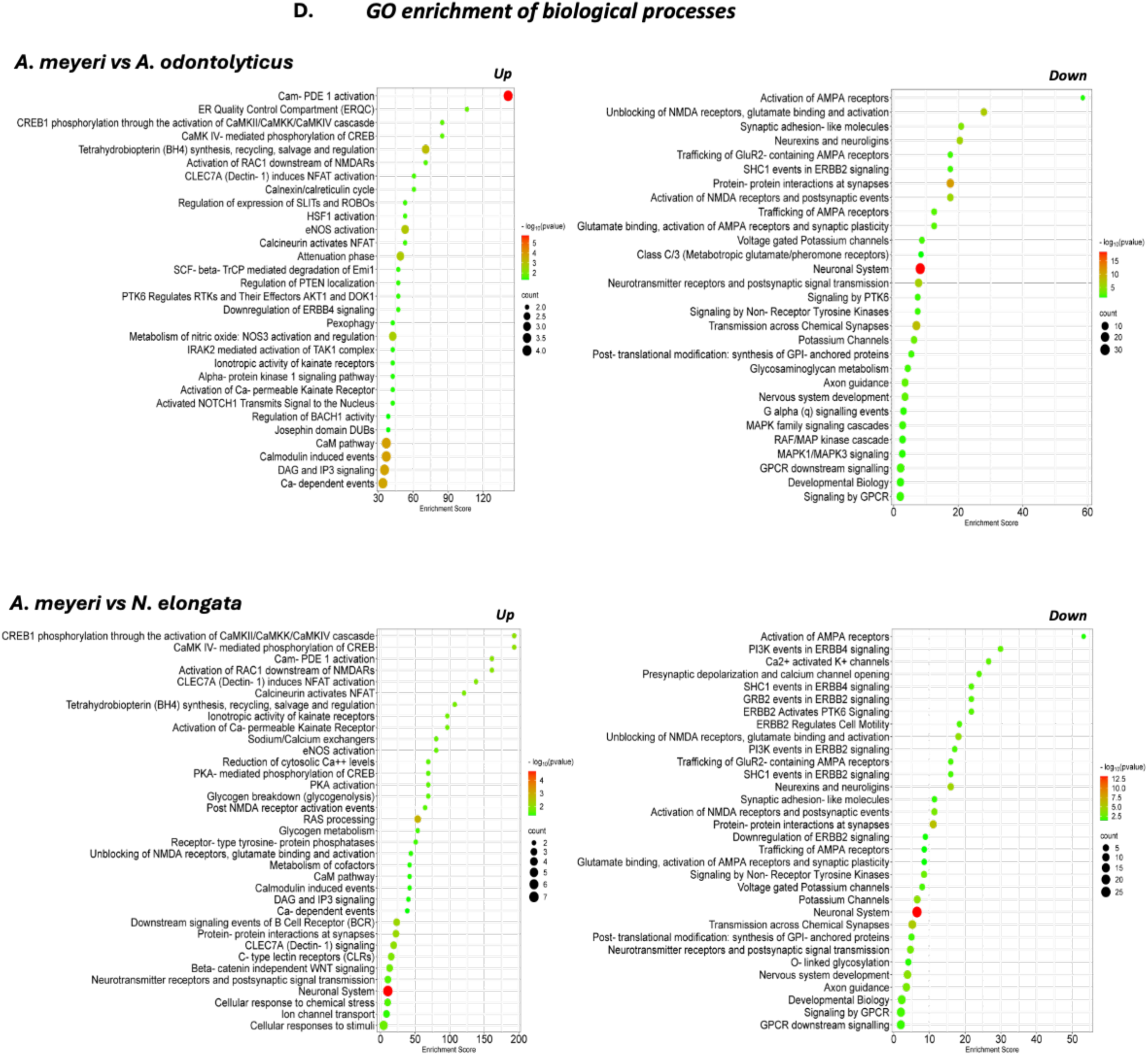
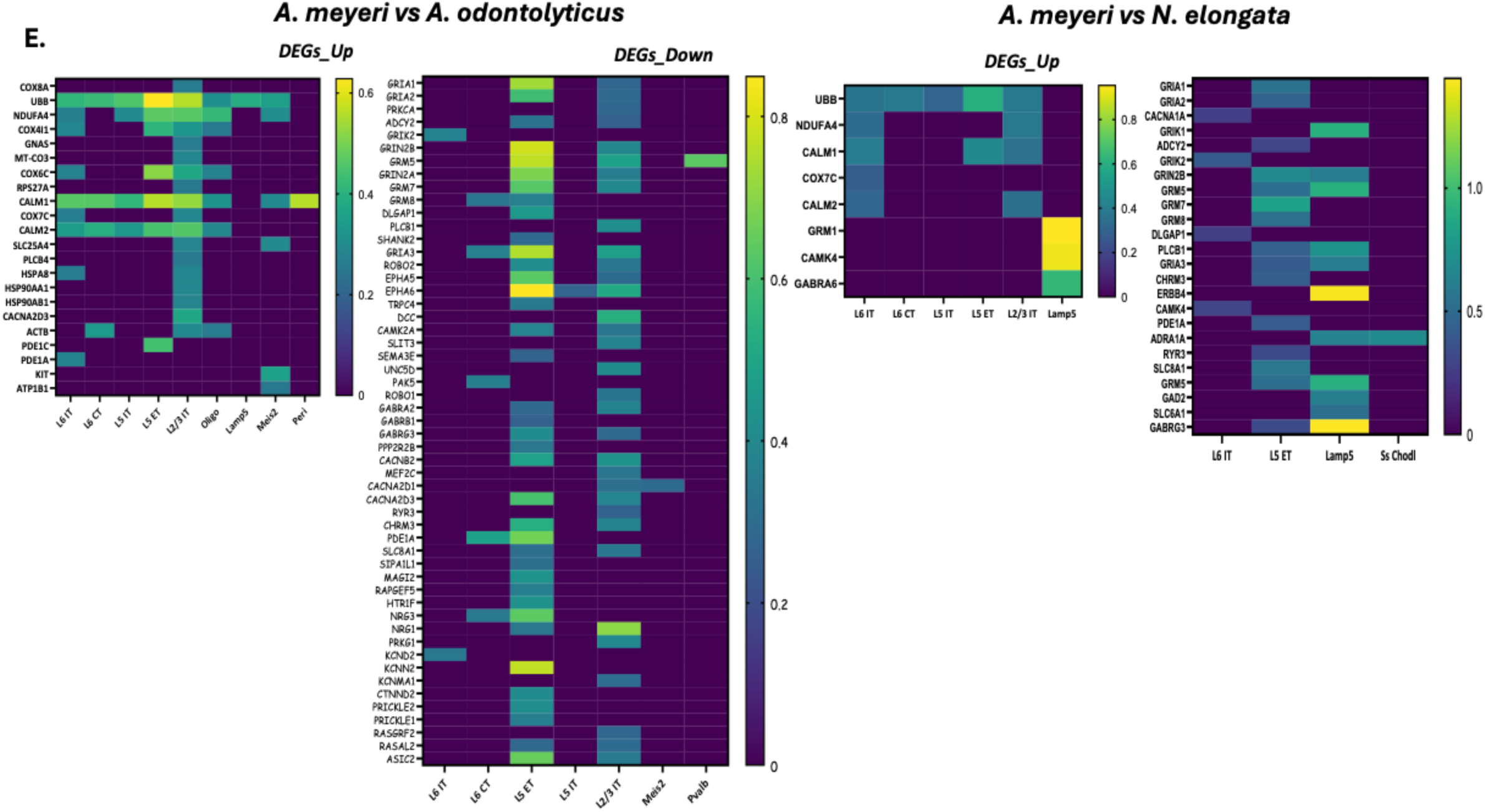
Cell counts along with the A. odontolyticus and N. elongata comparison with the A. meyeri group for scRNASeq analysis. **(a)** The single cells obtained from each integrated sample, and their UMAP plots, labeled with the cell types, showing that more excitatory neurons were detected and colored by condition. **(b)** The UMAP plots show the integrated dataset split by individual treatment as compared to the other microbiome groups. Twenty-two distinct cell types were identified through single-cell RNA sequencing, shown in different colors. **(c)** Gene ontology analysis of KEGG pathways representing the up-(pink) and down-regulated (blue) pathways, gene count (circle size), and their log10 p values in each dataset from different comparisons. **(d)** Gene ontology analysis based on Reactome/biological processes representing up- and down-regulated processes, -log(10) p-values (color coded), gene count (circle size), and their enrichment scores [-log10(geometric mean of p-values] through the DAVID bioinformatics functional annotation tool. **(f)** Heatmaps representing the up- and downregulated differentially expressed genes (according to the average log2 fold-change) in different cell types in each integrated dataset.

**Supplementary Fig. S5:**
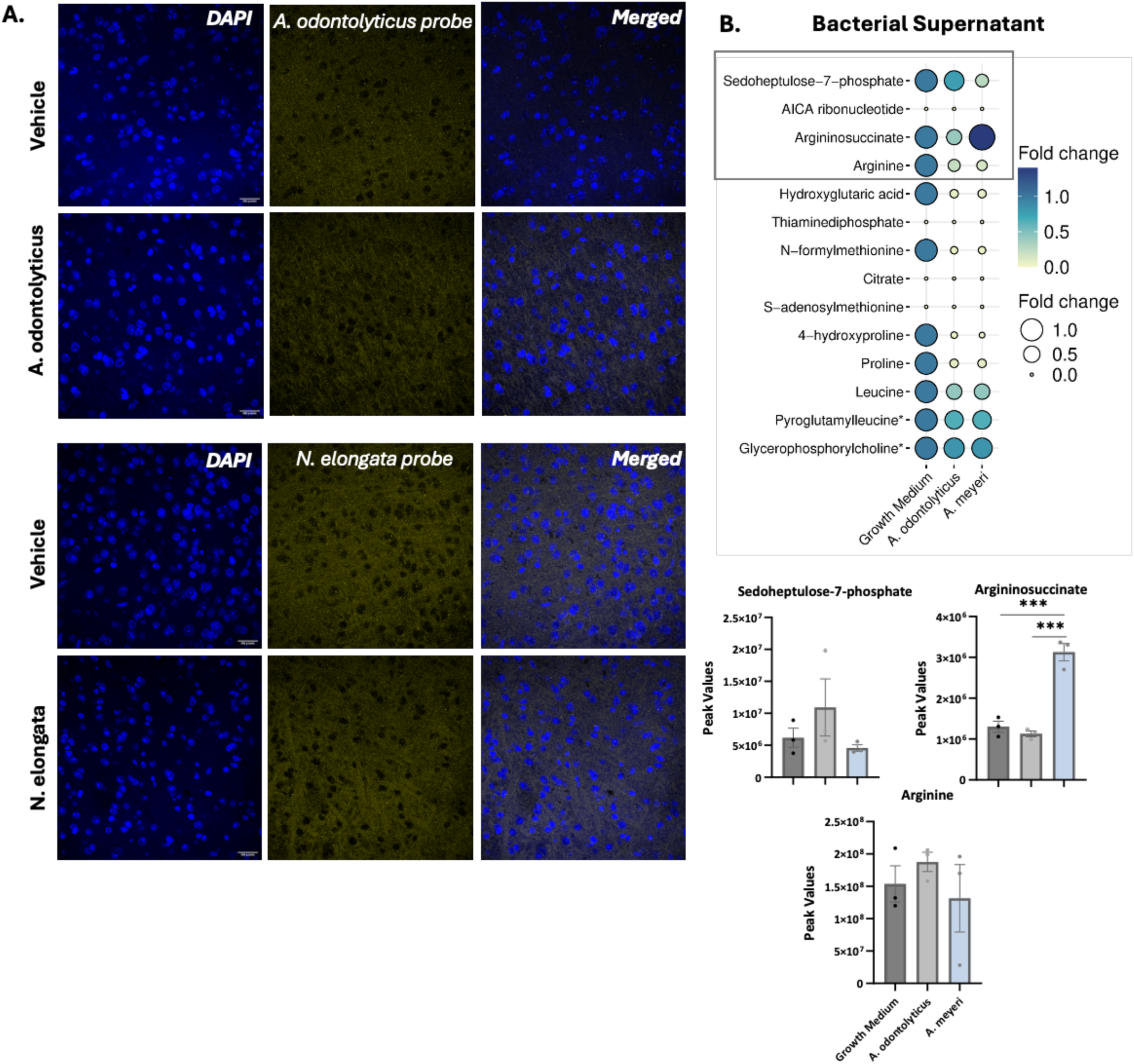
*A. odontolyticus* and *N. elongata genomes* were not detected in the brains by RNA-based insitu hybridization. **(a)** Representative images of 5µm FFPE sections from sagittal mouse brain sections were used for the RNAScope’s in situ hybridization protocol. The fluorescent assay shows no translocation of *A. odontolyticus* and *N. elongata* genomes from the oral cavity to the brain (yellow channel; Cy5), and DAPI is in blue. RNAScope Positive probe (UBC (Ubiquitin C) was used as a positive control targeting a common housekeeping gene in the Cy5 channel (Image is included in main figure with *A. meyeri* brains in Fig. 4). Images were captured at 40x by using Zeiss LSM 510 Meta confocal microscope (1024 × 1024 frame; n=4). **(b)** Out of 14 metabolites that were compared between saliva and the brain, 10 were found in the bacterial supernatants. The data is normalized with the abundance of metabolites present in their basal growth medium and presented as fold changes. Argininosuccinate was detected in the supernatant of *A. meyeri* (n=3 per group) by UPLC-MS. Individual graphs are shown for the metabolites that were common in the mouse brain and oral swabs (n=3).

